# NSD2 is a requisite subunit of the AR/FOXA1 neo-enhanceosome in promoting prostate tumorigenesis

**DOI:** 10.1101/2024.02.22.581560

**Authors:** Abhijit Parolia, Sanjana Eyunni, Brijesh Kumar Verma, Eleanor Young, Lianchao Liu, James George, Shweta Aras, Chandan Kanta Das, Rahul Mannan, Reyaz ur Rasool, Jie Luo, Sandra E. Carson, Erick Mitchell-Velasquez, Yihan Liu, Lanbo Xiao, Prathibha R. Gajjala, Mustapha Jaber, Xiaoju Wang, Tongchen He, Yuanyuan Qiao, Matthew Pang, Yuping Zhang, Mohammed Alhusayan, Xuhong Cao, Omid Tavana, Caiyun Hou, Zhen Wang, Ke Ding, Arul M. Chinnaiyan, Irfan A. Asangani

## Abstract

The androgen receptor (AR) is a ligand-responsive transcription factor that binds at enhancers to drive terminal differentiation of the prostatic luminal epithelia. By contrast, in tumors originating from these cells, AR chromatin occupancy is extensively reprogrammed to drive hyper-proliferative, metastatic, or therapy-resistant phenotypes, the molecular mechanisms of which remain poorly understood. Here, we show that the tumor-specific enhancer circuitry of AR is critically reliant on the activity of Nuclear Receptor Binding SET Domain Protein 2 (NSD2), a histone 3 lysine 36 di-methyltransferase. NSD2 expression is abnormally gained in prostate cancer cells and its functional inhibition impairs AR trans-activation potential through partial off-loading from over 40,000 genomic sites, which is greater than 65% of the AR tumor cistrome. The NSD2-dependent AR sites distinctly harbor a chimeric AR-half motif juxtaposed to a FOXA1 element. Similar chimeric motifs of AR are absent at the NSD2-independent AR enhancers and instead contain the canonical palindromic motifs. Meta-analyses of AR cistromes from patient tumors uncovered chimeric AR motifs to exclusively participate in tumor-specific enhancer circuitries, with a minimal role in the physiological activity of AR. Accordingly, NSD2 inactivation attenuated hallmark cancer phenotypes that were fully reinstated upon exogenous NSD2 re-expression. Inactivation of NSD2 also engendered increased dependency on its paralog NSD1, which independently maintained AR and MYC hyper-transcriptional programs in cancer cells. Concordantly, a dual NSD1/2 PROTAC degrader, called LLC0150, was preferentially cytotoxic in AR-dependent prostate cancer as well as NSD2-altered hematologic malignancies. Altogether, we identify NSD2 as a novel subunit of the AR *neo*-enhanceosome that wires prostate cancer gene expression programs, positioning NSD1/2 as viable paralog co-targets in advanced prostate cancer.

## Main

Prostate cancer is the most commonly diagnosed malignancy in North American men, with over 95% of the primary disease expressing the androgen receptor (AR) protein^1^. AR is a transcription factor that dimerizes and shuttles into the nucleus upon binding to its ligand (i.e., androgen), where it activates the expression of genes that drive terminal differentiation of luminal epithelial cells. In concert with chromatin and epigenetic regulatory proteins, AR primarily binds at distal *cis*-regulatory sites (also known as enhancers) containing a canonical androgen response element (ARE) that comprises a 15-bp palindromic DNA sequence with two invertedly-oriented hexameric 5′-AGAACA-3′ half-sites^2^. These DNA half-sites are separately recognized by each half of the AR homodimer^3^.

In prostate cancer cells, AR activity is extensively reprogrammed to enable and maintain malignant phenotypes^4–6^. Consequently, the androgen/AR axis is the primary target of all therapies following surgical resection or radiation of the organ-confined disease^7^. This acute dependency on AR activity is further reinforced in relapsed metastatic castration-resistant prostate cancer (mCRPC) through activating mutations or copy amplification of AR or its cofactors^8–12^. Seminal studies profiling the AR cistrome in primary prostate cancer uncovered *de novo* genesis of enhancers in the malignant state (neo-enhancers), resulting in a 2-3 fold expansion of the AR enhancer circuitry^5,6,13–15^. This engenders an acute dependency on chromatin-binding AR cofactors, such as SWI/SNF, BRD4, MED1, and p300/CBP, all of which have been independently assessed for therapeutic druggability in mCRPC^16–22^. Yet, the molecular mechanisms underlying chromatin redistribution of AR upon transformation or distinctive subunits of the AR transcriptional complex that assembles at neo-enhancer elements (i.e., the neo-enhanceosome) remain entirely unknown and, thus, unexplored for therapeutic targetability.

In this study, using an epigenetics-targeted functional CRISPR screen, we identified NSD2 (also known as MMSET, WHSC1) as a novel subunit of the AR enhanceosome complex in prostate cancer cells. NSD2 is a histone 3 lysine 36 mono and di-methyltransferase that activates gene expression by protecting the chromatin from accumulating repressive epigenetic marks, such as H3K27me3^23–25^. NSD2 is a bonafide oncogene in hematologic cancers and harbors recurrent activating alterations in over 15-20% of multiple myeloma^26–28^ and 10% of childhood acute lymphoblastic leukemia^29–31^. Notably, NSD2 hotspot mutations in the catalytic SET domain (e.g., E1099K) are significantly enriched in the relapsed disease^32,33^, which leads to a global increase in H3K36me2 levels and oncogenic remodeling of the epigenome^29,34,35^.

In prostate cancers, we found NSD2 to be exclusively expressed in the transformed cells, with no detectable expression in the normal epithelia, where it directly interacts with AR to enable its binding at chimeric AR-half motifs in concert with FOXA1 or other driver oncogenes. Inactivation of NSD2 entirely disrupted AR binding at over 65% of its tumor cistrome, importantly without affecting AR protein levels, and attenuated all major hallmark cancer phenotypes. NSD2 deficiency also engendered an increased dependency on NSD1, positioning the two paralogs as a digenic dependency. Concordantly, a dual NSD1/2 PROTAC degrader called, LLC0150, showed selective potency in AR-dependent as well as NSD2-altered human cancers. These findings mechanistically explain how AR gets reprogrammed, away from pro-differentiation physiological functions, to instead fuel prostate cancer growth and survival, and offer NSD1 and NSD2 as novel therapeutic vulnerabilities in the advanced disease.

## Results

### Epigenetics-focused CRISPR screen identifies NSD2 as a novel AR *neo*-coactivator in prostate cancer cells

Conventional plasmid-based reporter systems fail to capture intricate epigenetic or chromatin-level regulation of gene expression as they lack the native histone composition or higher-order chromosomal structure. Thus, we engineered an endogenous AR reporter system by using the CRISPR/Cas9 and homologous recombination methodologies. We edited the *KLK3* gene locus in AR-driven LNCaP cells to precisely knock-in the mCherry coding sequence directly downstream of the endogenous promoter and fused in-frame via an endopeptidase sequence to the *KLK3* gene (**Fig. 1a and Fig. S1a-c**). In the monoclonal reporter cell line, akin to PSA, mCherry expression is directly regulated by the AR transcriptional complex (**Fig. S1d-f**) and, most importantly, even captures chromatin or epigenetic-level changes in AR transactivation potential. Consistent with prior reports, like PSA, mCherry expression was attenuated upon pharmacologic inhibition of co-activators like BRD4^16^, SWI/SNF^18^, or P300/CBP^19^ as well as increased upon inhibition of the repressive PRC2/EZH2 complex^36,37^ (**Fig. 1b and Fig. S1g,h**). We used these endogenous AR reporter cell lines to carry out a functional CRISPR screen, wherein cells were treated with a custom sgRNA library targeting druggable transcriptional cofactors ^38^ (over 1500 sgRNAs against 200 plus genes; **Fig. S1h**) for 8 days, stimulated with DHT for 16h, and FACS-sorted into mCherry^HIGH^ and mCherry^LOW^ populations. This was followed by deep sequencing of the genomic sgRNAs, and the ratio of normalized sgRNA counts in mCherry^LOW^ to mCherry^HIGH^ cell populations was used to rank the individual guides. Here, ranked alongside BRD4^16^ and TRIM24^39,40^, we identified NSD2 as a novel AR coactivator in LNCaP cells (**Fig. 1c**). In contrast, subunits of the PRC2 complex, namely EZH2 and JARID2, that represses AR activity^36^ were enriched in the mCherry^HIGH^ cells. NSD2 is a histone 3 lysine 36 mono and di-methyltransferase (H3K36me2) that is centrally linked to the promotion of active gene expression^31,34^. Validating the screening results, siRNA-mediated knockdown of NSD2 significantly attenuated the expression of KLK3 in unaltered prostate cancer cells (**Fig. 1d**).

**Figure 1:**
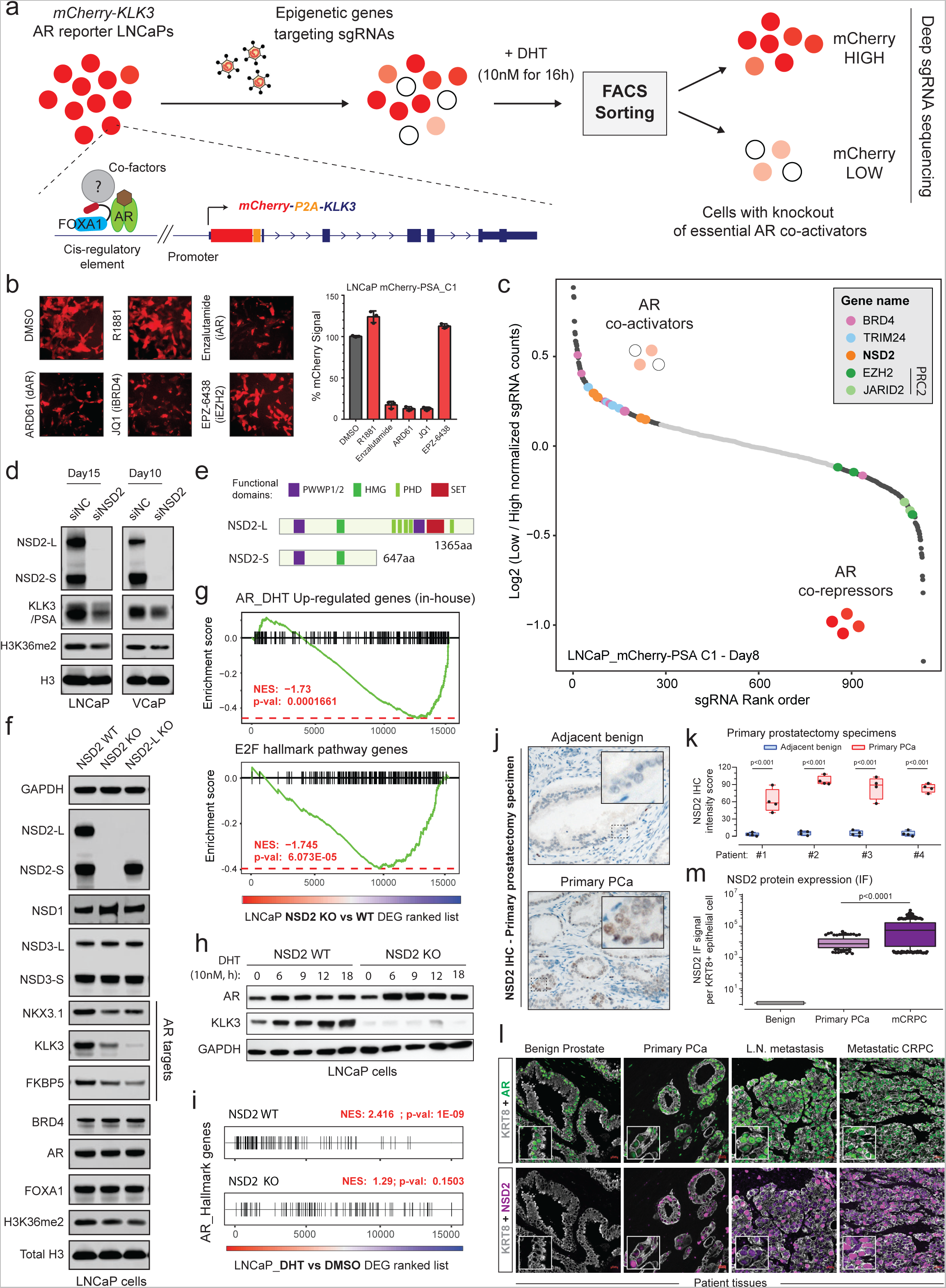
Epigenetics-CRISPR screen reveals NSD2 as a novel AR coactivator. a) Schematic of the epigenetic-targeted functional CRISPR screen using the LNCaP mCherry-KLK3 endogenous AR reporter lines. b) *Left*: mCherry immunofluorescence images of the reporter LNCaP cells treated with epigenetic drugs targeting known AR cofactors. *Right*: Barplot showing quantification of the mCherry signal from treated LNCaP reporter cells normalized to the DMSO treatment control. c) sgRNA enrichment rank plot based on the ratio of guide RNA abundances in mCherry-LOW to mCherry-HIGH cell populations. LNCaP reporter cells were treated with the sgRNA library for 8 days and FACS-sorted as shown in panel (a) for genomic sgRNA sequencing. d) Immunoblots of AR targets and histone marks upon treatment with the control (siNC) or NSD2-targeting (siNSD2) siRNAs in prostate cancer cell lines. Total histone H3 is used as the loading control. e) Representative protein map of NSD2-Long (NSD2-L) and NSD2-Short (NSD2-S) splice isoforms. HMG: High mobility group; PHD: Plant homeodomain. f) Immunoblots of AR targets and histone marks in CRISPR-mediated stable knock-out (KO) of both or only the NSD2-L isoform. Total H3 is the loading control. g) GSEA plots for AR and E2F up-regulated genes using the fold change rank-ordered genes from the LNCaP NSD2 knock-out (KO) vs wild-type (WT) control lines. DEGS, differentially expressed genes. (n=2 biological replicates) h) Immunoblots of labeled proteins in LNCaP NSD2 knock-out cells stimulated with 10nM DHT for increasing time durations. i) GSEA plots of AR hallmark genes in the LNCaP NSD2 wild-type (WT) vs knock-out (KO) cells using the fold change rank-ordered genes from DHT (10nM for 24h) vs DMSO treated LNCaP cells. DEGS, differentially expressed genes. j) Representative immunohistochemistry (IHC) images of NSD2 in primary prostatectomy patient specimens. k) NSD2 signal intensity plot from IHC staining in panel (j). Adjacent benign and primary prostate cancer tissues were used for this matched analysis. (n = 4 biological replicates; two-sided *t-test*). In box plots, the center line shows the median, box edges mark quartiles 1-3, and whiskers span quartiles 1-3 土1.5*interquartile range. l) Representative multiplex immunofluorescence (IF) images of KRT8, AR, and NSD2 in benign prostate, primary prostate cancer, or metastatic CRPC patient specimens. m) Quantification of NSD2 IF signal intensity per KRT8+ luminal epithelial cell from images in panel (l). (two-sided *t-test*). In box plots, the center line shows the median and the whiskers are drawn down to the 10th percentile and up to the 90th. Points below and above the whiskers are drawn as individual dots.

The NSD2 gene templates two splice isoforms producing a long, catalytically active form (hereafter referred to as NSD2-L) as well as a truncated shorter isoform (called NSD2-S) containing only the reader and protein-protein interaction PWWP and HMG domains, respectively—both of which are robustly expressed in PCa cells (**Fig. 1e and Fig. S2a**). Deletion of NSD2-L alone strongly attenuated the expression of AR target genes in LNCaP cells, which was comparable to the complete loss of the NSD2 protein (**Fig. 1f**), suggesting the methyltransferase function of NSD2 to be critical in sustaining AR activity. The transcriptomic analysis further revealed global AR activity to be significantly dampened in NSD2-deficient LNCaP cells with a parallel loss in hyper-proliferative gene expression programs (**Fig. 1g**). Notably, there was no loss in the abundance of the AR protein itself in NSD2-deleted cells (**Fig. 1f**), yet stimulation with DHT failed to significantly up-regulate the expression of cognate AR target genes (**Fig. 1h,i and Fig. S2b,c**).

To date, several studies have implicated NSD2 in prostate cancer^41–45^; however, it is worth noting that these studies were focused on metastatic, castration-resistant forms of the disease and not primary prostate tumorigenesis. Using tissue microarrays, these studies also described NSD2 protein expression to be elevated in cancer specimens, showing a stage-wise increase from primary to metastatic castration-resistant or transdifferentiated neuroendocrine prostate cancer^43,44^. Building on these findings, in primary prostatectomy specimens we strikingly found NSD2 levels to be undetectable in the normal or adjacent benign foci with a marked gain in expression in the malignant lesions (**Fig. 1j,k and Fig. S2f**). Consistent with this, using a single-cell RNA-seq dataset, akin to the cancer-specific *PCA3* transcript ^46^, we found the *NSD2* transcript to be expressed exclusively in the AR+ luminal epithelial cells derived from the tumor (**Fig. S2d**). Pseudo-bulk analyses of luminal epithelial cells from patients (n=18) confirmed NSD2 expression to be markedly elevated in the matched tumor vs the normal compartment, with NSD2 expression also positively correlating with Gleason score of the primary disease (**Fig. S2e,g**). Using a more sensitive and multiplex immunofluorescence approach, we further confirmed the KRT8+/AR+ normal epithelial cells to have no detectable expression of the NSD2 protein, which was exclusively and robustly expressed in the nuclei of transformed luminal epithelial cells with the highest expression in metastatic tumors (**Fig. 1l,m**). Altogether, this data suggests that NSD2 is abnormally expressed in the transformed primary prostate luminal epithelial cells, wherein its function is critical for maintaining the transactivation potential of the AR signaling complex, making NSD2 one of the first reported *neo*-coactivators of the AR transcriptional complex.

### NSD2 expands the AR enhancer circuitry to include chimeric AR-half motifs in prostate cancer

Given that NSD2 loss had no impact on the abundance of the AR protein, we next profiled AR binding on chromatin. Here, ChIP-seq of AR in NSD2-deficient LNCaP cells uncovered a dramatic and complete off-loading of the AR protein from over 40,000 genomic sites or over 65% of its tumor cistrome (**Fig. 2a**). The majority of the lost sites (i.e., NSD2-dependent) were within intronic or distal intergenic regions that are associated with *cis*-regulatory enhancer elements (**Fig. 2b**). Intriguingly, at these sites there was no change in the binding of FOXA1 upon inactivation of NSD2. Yet, the loss of chromatin binding of AR was sufficient to trigger loss of the H3K27Ac mark that demarcates active enhancers (**Fig. 2c**). In contrast, AR continued to robustly bind at over 20,000 genomic elements independent of NSD2, which also retained the H3K27Ac active enhancer mark in the NSD2-null prostate cancer cells (**Fig 2a-c**).

**Figure 2:**
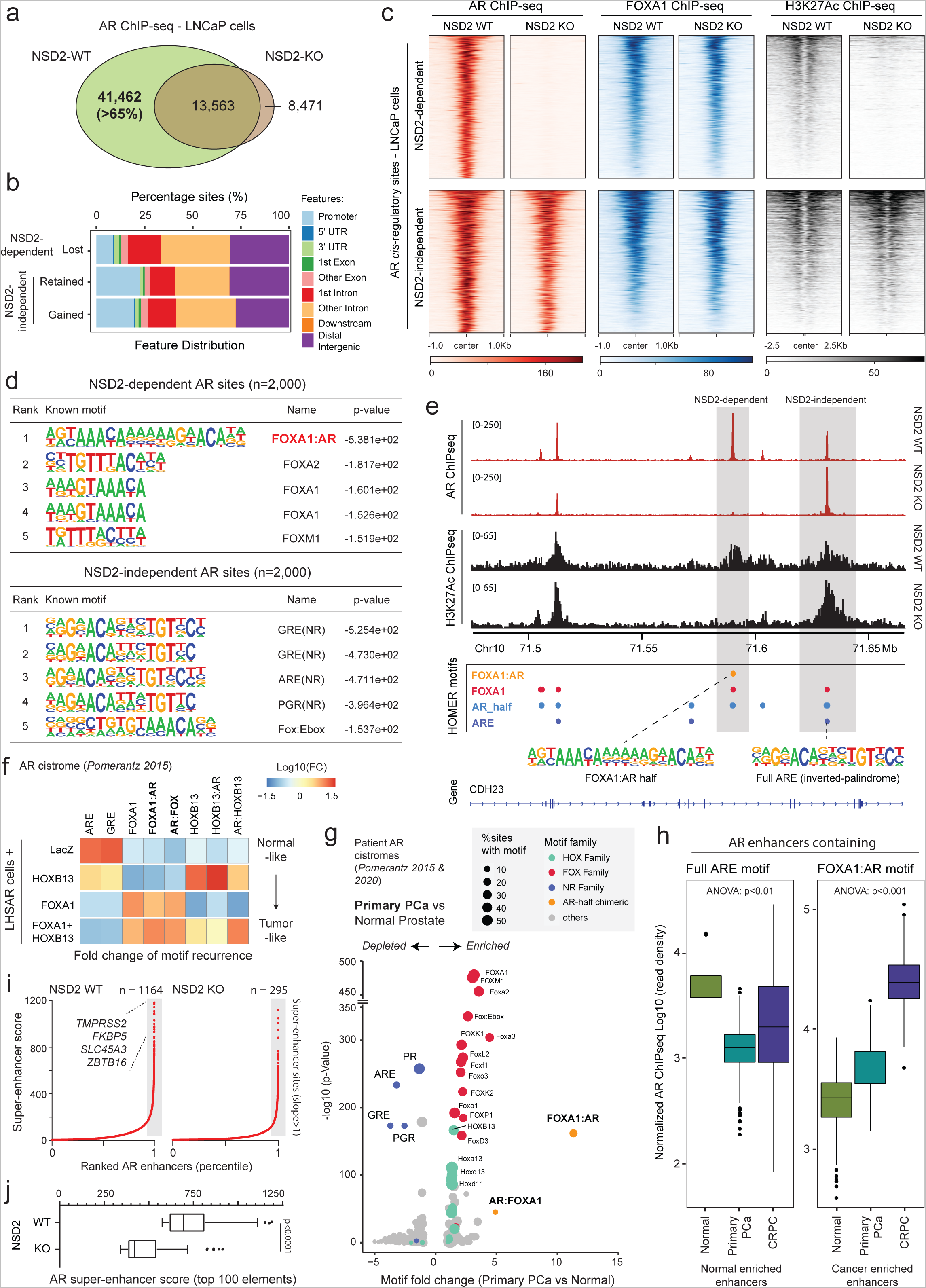
NSD2 expands the AR neo-enhancer circuitry to include chimeric AR half-sites. a) Venn diagram showing overlaps of AR ChIP-seq peaks in NSD2 wild-type (WT) and knock-out (KO) LNCaP cell lines. b) Genomic location of NSD2-dependent and independent AR sites defined from the overlap analysis in panel (a). c) ChIP-seq read-density heatmaps of AR, FOXA1, and H3K27Ac at top 1,000 AR enhancer sites (ranked by score) in LNCaP NSD2 WT and KO cell lines. d) Top five known HOMER motifs (ranked by p-value) enriched within NSD2-dependent and independent AR sites in LNCaP cells. (HOMER, hypergeometric test). e) ChIP-seq read-density tracks of AR and H3K27Ac within a Chr10 locus in NSD2 WT and KO LNCaP cell lines. HOMER motifs detected within AR peaks are shown below with grey boxes highlighting NSD2 dependent and independent AR elements. f) Fold change heatmap of HOMER motifs enrichment within AR binding sites specific to HOXB13, FOXA1, or FOXA1+HOXB13 overexpression in LHSAR cells (data from Pomerantz et al, 2015). g) Fold change and significance of HOMER motifs enriched within primary prostate cancer-specific AR sites over normal tissue-specific AR elements (data from Pomerantz et al., 2015). h) AR ChIP-seq read density boxplot at sites containing the ARE or the FOXA1:AR chimeric motif in primary normal and tumor patient samples (normal prostate, n=7; primary prostate cancer, n=13; CRPC, n=15). In box plots, the center line shows the median, box edges mark quartiles 1-3, and whiskers span quartiles 1-3 土 1.5 * interquartile range. i) Rank-ordered plot of AR super-enhancers (HOMER ROSE algorithm) in NSD2 wild-type and knock-out LNCaP cells with select *cis*-coded known AR target genes noted. j) Boxplot of AR super-enhancer scores (HOMER ROSE al) of top 100 *cis*-elements in NSD2 WT or KO LNCaP cells.

Motif analyses (HOMER^47^) of the NSD2-dependent AR sites revealed a chimeric motif comprising a FOXA1 element juxtaposed to the AR-half site (annotated as the FOXA1:AR-half motif) as the most significantly enriched DNA sequence within 200bp centered at the binding peak (**Fig. 2d**). Almost 40% of the NSD2-dependent AR enhancers harbored the FOXA1:AR-half motif (**Table S1 and S2**). In contrast, NSD2-independent AR sites (i.e., ~35% of the tumor cistrome), a large fraction of which showed increased AR binding upon NSD2 inactivation, housed the canonical androgen response element (ARE) comprised of a 15-bp palindromic sequence with two invertedly-oriented hexameric 5′-AGAACA-3′ half-sites (**Fig. 2d**). The two modes of AR chromatin interaction were strikingly evident within a gene locus on Chr10, wherein the loss of NSD2 led to a complete disruption of AR binding at the site containing the FOXA1:AR-half motif, without affecting its binding at canonical AREs in the immediate *cis*-neighborhood, with concordant changes in the H3K27Ac signal (**Fig. 2e**). Furthermore, custom motif analyses revealed an enrichment of motifs of other oncogenic transcription factors, including HOXB13 and ETS, within 25bp of the AR-half element at the NSD2-dependent AR sites (**Fig. S3a**). Next, we custom assembled chimeric AR-half motifs with FOXA1 and HOXB13 elements (in both 5’ and 3’ confirmations, see Methods) and interrogated their recurrence in a published isogenic AR ChIP-seq dataset derived from the non-malignant immortalized LHSAR cells^5^. Here, we found the overexpression of FOXA1 and HOXB13 alone or in combination in LHSAR cells to markedly shift the AR cistrome away from full AREs (normal-like) toward the chimeric AR-half elements in the tumor-like state (**Fig. 2f and Fig. S3b**). Most strikingly, motif analysis of the AR cistromes generated from primary patient specimens^5,6^ revealed the FOXA1:AR-half motif to be exclusively detected in the tumor-specific AR enhancer circuitries, with such chimeric motifs being essentially absent within the normal tissue-specific AR sites (**Fig. 2g and Fig. S3c-e**). In the same analyses, we found the palindromic AREs to be depleted within cancer-specific enhancers compared to AR sites specific to the normal prostate (**Fig. 2g**). Accordingly, AR ChIP-seq signal at elements comprising full AREs showed the strongest AR binding in the normal tissues, while enhancers housing the chimeric FOXA1:AR-half motif had significantly higher AR occupancy in cancer tissues (**Fig. 2h**). H3K27Ac ChIP-seq signal from matched patient specimens showed a similar reprogramming, with the FOXA1:AR-half sites being strongly activated in mCRPC tumors (**Fig. S3f**).

In tumor cells, the aberrant expression of driver oncogenes is frequently amplified through dense clusters of closely spaced enhancers, often referred to as super-enhancers^48^. Notably, NSD2 inactivation also resulted in the loss of over 75% of the AR super-enhancers (**Fig. 2i**), including those that are hijacked by activating translocations in the *TMPRSS2* and *SLC45A3* gene loci^49^ (**Fig. S3g**). The residual super-enhancer elements also showed a marked decrease in transcriptional activity in the NSD2 knock-out relative to the wild-type LNCaP cells (**Fig. 2j**). Altogether, these data suggest that, upon ectopic expression in prostate cancer cells, NSD2 critically assists oncogenic transcription factors (such as FOXA1 and HOXB13) in expanding the AR enhancer circuitry to include chimeric AR-half sites that constitute over two-thirds of the malignant cistromes. In other words, NSD2 plays a critical role in loading the AR enhanceosome at cancer-specific, degenerate AR-half elements that lead to a 2-3 fold expansion of the AR *neo*-enhancer circuitries.

### NSD2 enables oncogenic AR activity with NSD1/2 emerging as paralog co-dependencies

Given the loss of NSD2 function resulted in targeted disruption of the cancer-specific AR cistrome, we set out to phenotypically characterize the NSD2-deficient AR-positive prostate cancer cells. Here, siRNA knockdown or CRISPR knockout of NSD2 significantly impaired the hyper-proliferative ability of several AR+ prostate cancer cells (**Fig. 3a and Fig. S4a**). NSD2-deficient cells also entirely lost the ability to invade through Matrigel in the Boyden chamber assay (**Fig. 3b**), as well as form colonies from a single cell in clonogenic assays (**Fig. 3c and Fig. S4b**). 22RV1 cells lacking NSD2 also had a dramatic attenuation of the ability to form xenografts when injected subcutaneously into the dorsal flanks of NOD/SCID mice (**Fig. 3d**). Most strikingly, exogenous reintroduction of the catalytically competent NSD2-L restored the grafting ability in mice (green line, **Fig. 3d**), with the resulting tumors growing at a rate comparable to those established with the parental 22RV1 cells (**Fig. S4c**). NSD2-L re-expression also restored the invasive ability of NSD2 KO LNCaP cells (**Fig. S4d**) along with restoring the expression of KLK3 (**Fig. S4e**). In the same NSD2-deficient models, re-expression of NSD2 variant lacking the SET domain (dSET) failed to rescue KLK3 (**Fig. S4f**), while expression of the hyper-active NSD2-E1099K SET-domain mutant^29^ completely restored its levels (**Fig. S4g**). Next, we engineered the NSD2-null 22RV1 cells to stably express the exogenous dTAG-version of the NSD2-L protein fused to the FKBP12^F36V^ tag^50^. In these cells, NSD2-L-FKBP12^F36V^ could be rapidly degraded upon treatment with the dTAG-13 degrader of FKBP12^F36V^ (**Fig. S4h**). The 22RV1 NSD2 KO + NSD2-L-FKBP12^F36V^ cells successfully grafted and robustly grew to form tumors *in vivo*; however, dosing of host animals with the FKBP12^F36V^ PROTAC, dTAGv-1, significantly diminished the growth of tumor xenografts vs the control treatment (**Fig. 3e and Fig. S4i**). Even at the molecular level, degradation of the exogenous NSD2-L-FKBP12^F36V^ protein in NSD2-null LNCaP cells resulted in lower levels of KLK3 and the chromatin-associated phosphorylated (p-S81) form of AR without a decrease in total AR expression (top panel, **Fig. 3f**). Concordantly, chromatin fractionation in these cells showed a marked loss of chromatin-bound AR and the H3K36me2 mark upon degradation of NSD2-L-FKBP12^F36V^ with dTAG-13 (bottom panel, **Fig. 3f**). In the LNCaP NSD2-dTAG models, degradation of exogenous NSD2 also lead to downregulation of multiple AR target gene expression in a time-dependent manner (**Fig. S4j**). This data positions NSD2 as a molecular “switch” that wires the oncogenic AR cistrome, enabling the regulation of hallmark cancer phenotypes including hyper-proliferation, invasion, anchorage independence, as well as the grafting potential in mice.

**Figure 3:**
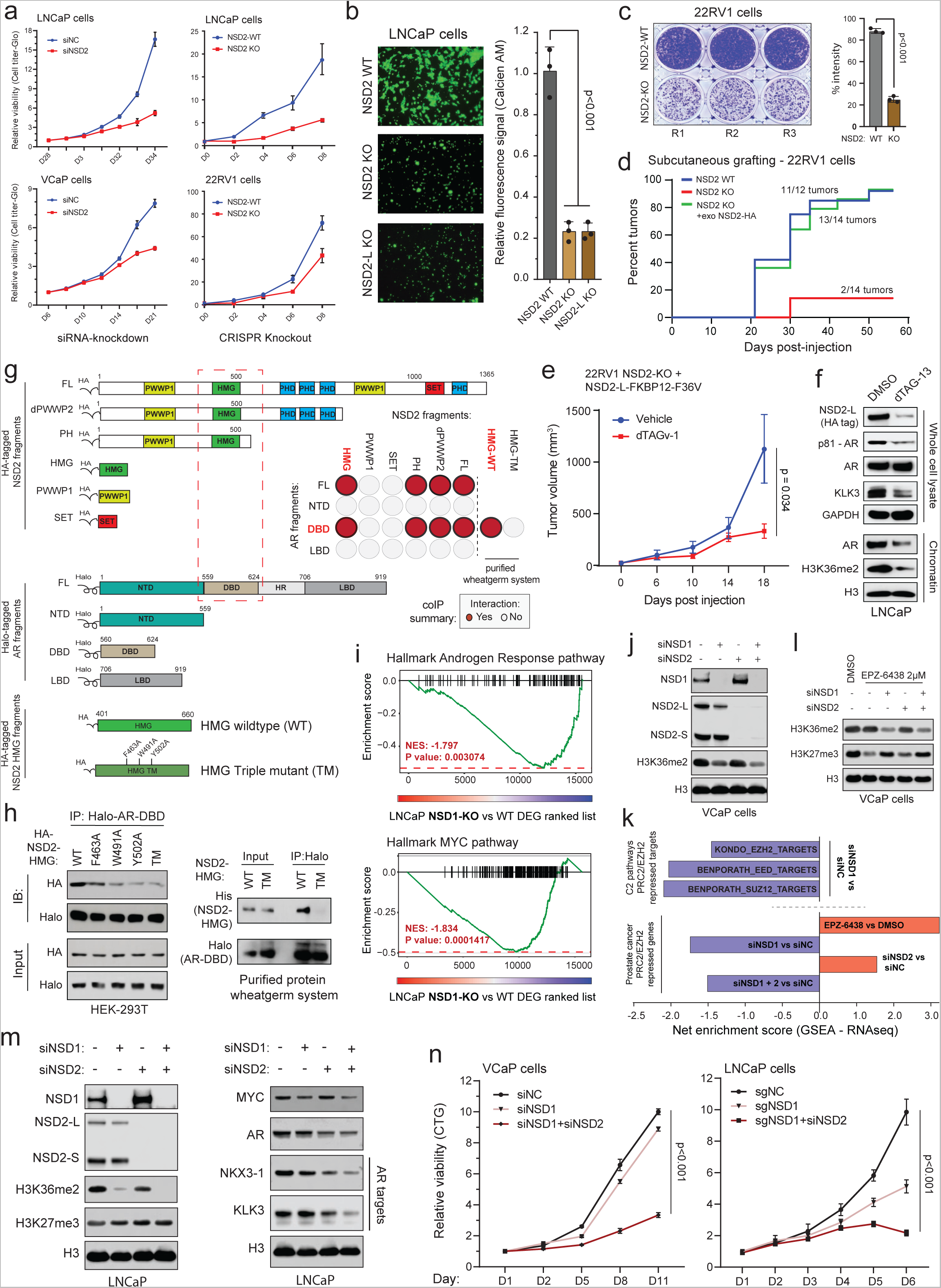
NSD2 enables oncogenic AR activity with NSD1/2 being paralog co-dependencies. a) *Left*: Growth curves (Cell-titer Glo) of prostate cancer cells treated with non-targeting (siNC) or NSD2-targeting siRNAs. *Right*: Growth curves (Cell-titer Glo) of CRISPR-engineered prostate cancer cells with deletion of the NSD2 gene (NSD2 KO) relative to the control wildtype (NSD2 WT) cells. b) *Left*: Representative images of Boyden chambers showing invaded cells stained with calcein AM dye in the LNCaP NSD2 WT and KO. *Right*: Barplot showing quantified fluorescence signal from invaded cells (two-way ANOVA and Tukey’s test) c) *Left*: Representative images of colonies of 22RV1 NSD2 WT and KO cell lines (n= 3 biological replicates). *Right*: Barplot showing staining intensity of the colonies. (Two-sided t-test) d) Reverse Kaplan Meier plot of subcutaneous tumor grafting of 22RV1 NSD2 WT, NSD2 KO, or NSD2 KO cells rescued with the NSD2-L-HA isoform. e) Tumor volumes of 22RV1 NSD2-KO+NSD2-FKBP12^F36V^ cell line-derived xenografts with or without treatment with dTAGv-1. Mean 土 SEM is shown. (Multiple t-test) f) Immunoblots of noted proteins in whole-cell or chromatin fractions of LNCaP NSD2-FKBP12^F36V^ cell line plus/minus treatment with dTAG-13 (0.5uM for 24h). g) Schematic of the epitope-tagged NSD2 and AR fragments used in the interaction studies. The dashed red box marks NSD2 and AR interacting functional domains. *Inset*: Summary of co-immunoprecipitation assays showing interaction between NSD2 and AR protein fragments. Source immunoblots are included in Fig. S5b,c. Red circles show interaction while grey circles represent no detectable binding between corresponding fragments. h) *Left*: Immunoblots of Halo-tag-based co-immunoprecipitation of the AR DNA-binding domain (DBD) in HEK293FT cells that overexpress HA-tagged NSD2-HMG single or triple mutants. Immunoblots of input fractions are shown as control. DBD: DNA binding domain; NSD2 HMG domain mutations: F463A, W491A, Y502A and, the triple mutant (TM). *Right*: Immunoblots of co-immunoprecipitation of wheatgerm-purified Halo-AR-DBD fragment with the purified His-NSD2 HMG variants. Immunoblots of input fractions are shown as control. i) GSEA plots for AR and MYC target genes using the fold change rank-ordered genes (RNAseq) from the LNCaP NSD1 KO vs wild-type cells. DEGS, differentially expressed genes. (n=2 biological replicates) j) Immunoblots of labeled proteins in VCaP cells upon siRNA treatment targeting NSD1 and/or NSD2 genes. Total histone H3 is a loading control. k) *Top*: GSEA net enrichment scores of EZH2/PRC2 repressed gene signatures (C2 pathways) in siNSD1 versus siNC treated VCaP cells. *Bottom*: GSEA net enrichment score of the prostate cancer-specific EZH2 repressed gene signature (defined in-house, see Methods) in siNSD1 and/or siNSD2 vs siNC treated VCaP cells. l) Immunoblots of noted histone marks in VCaP cells co-treated with siRNA targeting NSD1 and/or NSD2, followed by the EZH2-inhibitor EPZ-6438. m) Immunoblot of listed proteins in LNCaP cells treated with control siRNA (siNC) or siRNA targeting NSD1 and/or NSD2. n) *Left*: Cell growth curve (Cell-titer Glo) assays of VCaP cells treated with control siRNA (siNC) or siRNA targeting NSD1 or NSD1+NSD2 together. *Right*: Growth curves of NSD1-deficient (sgNSD1) LNCaP cells plus/minus treatment with siRNA targeting NSD2. (Two-way ANOVA or Tukey’s test).

In line with the above model, NSD2 co-eluted with higher order AR transcriptional complexes in size exclusion chromatography (**Fig. S4k**) as well as co-precipitated with AR in several prostate cancer cell lines (**Fig. S5a**). Using a fragment-based co-immunoprecipitation approach, we further mapped the high mobility group box (HMG-box) domain of NSD2 to interact with the DNA binding domain (DBD) of AR (**Fig. 3g and Fig. S5b,c**). Furthermore, alanine substitution of three highly conserved residues within the HMG-box domain (i.e., F436/W491/Y502A) alone or together (labeled as TM: triple mutant, **Fig. 3g**) disrupted its interaction with the AR-DBD in mammalian HEK293T ectopic system as well as eukaryotic cell-free wheatgerm extract purified system (**Fig. 3h**). This suggests that NSD2 directly, and independent of DNA, interacts with the AR DBD through its HMG-box domain, which is notably absent in other members of the NSD family of histone methyltransferases.

Despite a striking loss of defining neoplastic features, NSD2-deficient prostate cancer cells remained viable. Thus, we speculated if other NSD family paralogs could sustain AR activity through alternative mechanisms in the absence of NSD2. To test this, we CRISPR-engineered LNCaP cells to knockout NSD1 or NSD3 individually and assessed its impact on global transcription. Unlike NSD3, here we found NSD1 loss to significantly attenuate the hyper-transcribed AR and MYC gene programs (**Fig. 3i and Fig. S5d,e**), in addition to reducing the AR protein levels (**Fig. S5f**). NSD1 inactivation also resulted in a dampening of the hyper-proliferative gene pathways (like E2F and G2M; **Fig. S5g**) and had the strongest reduction in H3K36me2 levels upon a single-gene loss in prostate cancer cells (**Fig. 3j and Fig. S5f**). This positions NSD1 as the predominant H3K36 di-methyltransferase in prostate cancer cells. Using structural methods, the NSD-catalyzed H3K36me2 mark was shown to sterically hinder loading of the H3K27 residue into the catalytic pocket of the EZH2 enzyme^51^. Recent studies also reported NSD1 to be the dominant enzyme that antagonizes the repressive PRC2/EZH2 complex^52^, which is hyperactivated in prostate cancer cells and represses AR activity^36,37,53^. In line with these studies, we found NSD1 loss to lead to a significant increase in repressive activity of the EZH2/PRC2 complex, with several target gene signatures being strongly downregulated in NSD1 null versus wildtype VCaP cells (top panel, **Fig. 3k**). This was confirmed using a prostate cancer-specific PRC2 repressive gene signature, experimentally defined via treatment with a clinical-grade EZH2 inhibitor EPZ-6438 (see Methods), which was significantly downregulated upon loss of NSD1, but not NSD2, in VCaP cells (bottom panel, **Fig. 3k**). Concordantly, treatment of NSD1-deficient cells with EPZ-6438 still had substantially higher residual levels of the EZH2-catalyzed H3K27me3 repressive mark relative to the control as well as NSD2-deficient VCaP cells (**Fig. 3l**). In these experiments, loss of NSD2 alone had little to no effect on EZH2/PRC2 activity with the combined inactivation of NSD1 and NSD2 transcriptionally phenocopying the loss of NSD1 alone (**Fig. 3j-l**). These results position NSD1 as the primary writer of the H3K36me2 histone mark that counterbalances the EZH2/PRC2 repressive complex in prostate cancer cells to maintain the hyper-transcriptional AR and MYC gene programs.

Interestingly, the loss of NSD2 led to a significant increase in NSD1 levels in prostate cancer cells (**Fig. 3j,m**), likely suggesting that NSD1 could sustain residual AR activity in these cells, thus keeping them alive. Parallel inactivation of NSD1 and NSD2 in LNCaP and VCaP cells resulted in the strongest decrease in H3K36me2 levels and AR target gene expression (**Fig. 3j,m**), as well as resulted in an accumulation of apoptotic marker cleaved-PARP (**Fig. S5h**). Consistently, while NSD1 or NSD2 inactivation alone only attenuated the growth of prostate cancer cells, their combined loss resulted in significant cytotoxicity (**Fig. 3n**). Even analyses of the CRISPR DepMap data from Project Achilles^54,55^ showed either NSD1 or NSD2 deletion had no cytotoxic effect in prostatic cell lines (**Fig. S5i**). Altogether, these data suggest that NSD1 and NSD2, through distinct mechanisms, promote a hyper-transcriptional chromatin state and maintain oncogenic AR activity, respectively, in prostate cancer cells.

### LLC0150, a dual NSD1/2 PROTAC degrader, shows preferential cytotoxicity in AR/FOXA1-driven prostate cancer

Following a medicinal chemistry campaign, we developed a proteolysis targeting chimera (PROTAC) compound, called LLC0150, which co-targets NSD1 and NSD2 (**Fig. 4a**). LLC0150 links a recently published NSD2 PWWP-domain binding warhead^56^ to a cereblon E3-ligase-recruiting moiety pomalidomide. Treatment with LLC0150 led to the degradation of both NSD1 and NSD2, while completely sparing NSD3 (**Fig. 4b**), in a proteasome and cereblon-dependent manner (**Fig. S6a**). Global proteomics confirmed LLC0150 as a specific dual NSD1/2 degrader with no effect on other PWWP-domain-containing proteins (**Fig. S6b**). Consistent with our genetic approach, in prostate cancer cells, NSD1/2 co-degradation using LLC0150 triggered a striking decrease in the expression of AR and MYC, as well as their downstream gene targets (**Fig. 4c,d**). Acute loss of NSD1/2 in LNCaP cells treated with LLC0150 resulted in significantly diminished binding of AR and FOXA1 to chromatin (**Fig. 4e,f and Fig. S6c**), with a parallel loss of H3K27Ac activation mark at the core AR/FOXA1 enhancer elements (**Fig. 4f**). Treatment with LLC0150 led to a significant decrease in p-S81 AR levels and diminished loading of AR on chromatin that carried a significantly lower amount of the H3K36me2 chemical mark (**Fig. S6d**). Consequently, in the absence of NSD1/2, DHT-induced expression of AR target genes was significantly attenuated (**Fig. S6e**). LLC0150 treatment also markedly disrupted the assembly and activity of AR super-enhancers in LNCaP cells (**Fig. S6f**)

**Figure 4:**
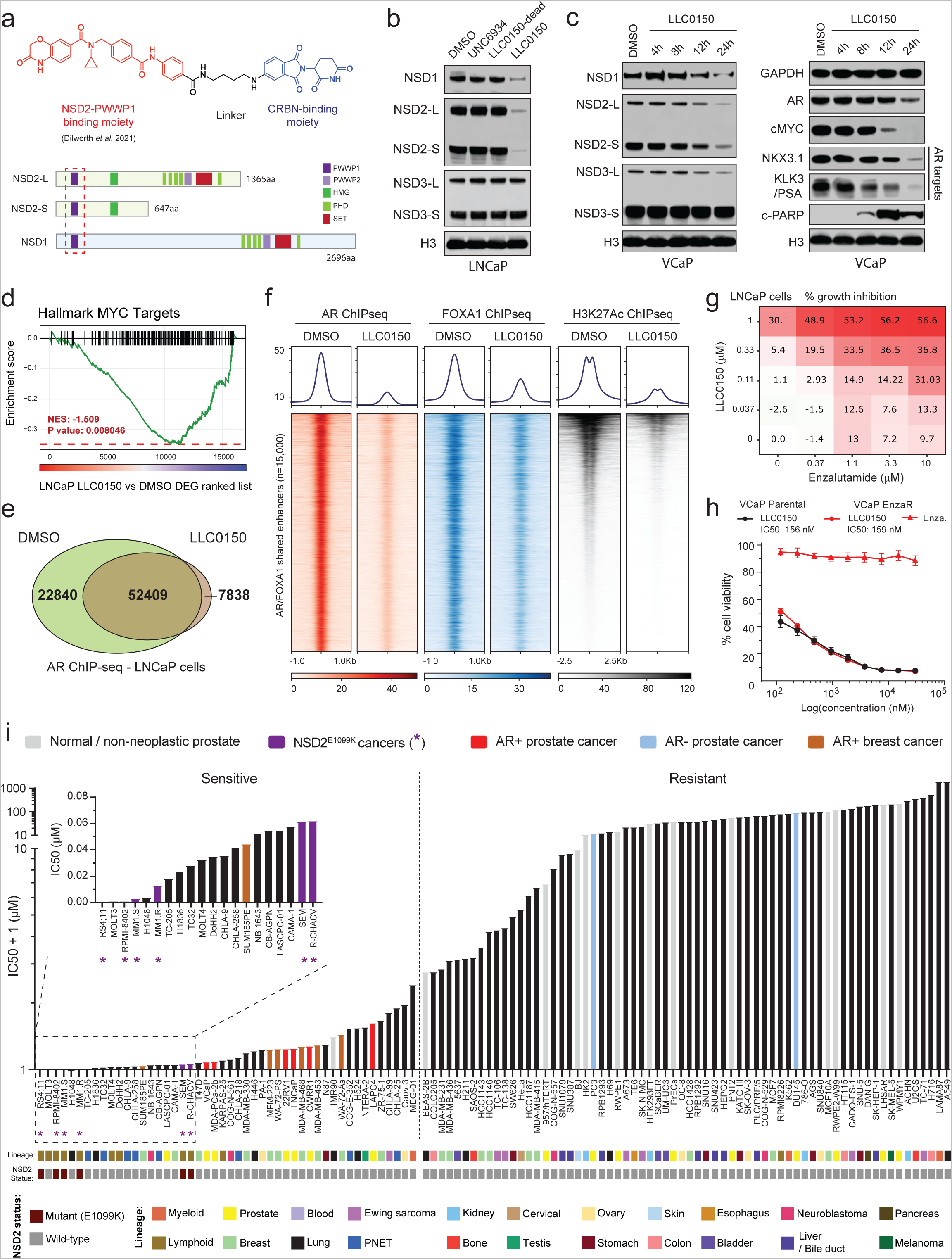
LLC0150 is an NSD1/2 PROTAC with preferential cytotoxicity in AR-driven prostate cancer. a) Structure of LLC0150 and schema of NSD1 and NSD2 functional domains. LLC0150-binding PWWP1 domain is highlighted using a dashed red box. HMG: High mobility group; PHD: Plant homeodomain. b) Immunoblots of listed proteins in VCaP cells treated with UNC6934 (warhead), LLC0150-dead (epimer control) or LLC0150 for 12h at 1uM. Total histone H3 is used as a loading control. c) Immunoblots of listed proteins in VCaP cells treated with LLC0150 (2uM) for increasing time durations. Total histone H3 is used as a loading control. d) GSEA plots of MYC target genes using the fold change rank-ordered genes from LLC0150 vs DMSO treated LNCaP cells. DEGS, differentially expressed genes. e) Venn diagram showing the overlap of AR ChIP-seq peaks in LNCaP cells treated with LLC0150 (2uM for 48h) or DMSO as control. f) ChIP-seq read-density heatmaps of AR, FOXA1, and H3K27Ac at enhancers that are co-bound by AR and FOXA1 in LNCaP cells plus/minus treatment with LLC0150 (2uM for 48h). g) Percent growth inhibition (Cell-titer Glo) of LNCaP cells upon co-treatment with varying concentrations of LLC0150 and enzalutamide. h) Dose-response curves of LLC0150 or enzalutamide in parental or enzalutamide-resistant VCaP cells. Data are presented as mean +/− SEM (n=2 biological replicates). Serving as a control, enzalutamide dose-response curve credentials the enzalutamide-resistant VCaP cell line. i) IC50 rank-order plot of over 110 human-derived normal or cancer cell lines after 5 days of treatment with LLC0150. AR+ prostate cancer models are highlighted in red, and NSD2-mutant hematologic cell lines are shown in purple as well as marked with an asterisk (*). Each cell line’s originating tissue lineages and known NSD2 alteration status are shown below.

In line with disruption of the AR/FOXA1 neo-enhancer circuitry, global transcriptomic analyses of LNCaP and VCaP cells treated with LLC0150 showed a significant attenuation of proliferative pathways with a parallel induction of apoptotic cell death signaling (**Fig. S7a**). This was confirmed via massive accumulation of cleaved-PARP in the LLC0150-treated AR-positive prostate cancer cell lines (**Fig. 4c and Fig. S7b**). AR-positive prostate cancer cell lines were considerably more sensitive to treatment with LLC0150 relative to AR-negative cancer, immortalized normal, as well as primary prostate epithelial cells (VCaP: 84.8 nM, LNCaP: 251.6 nM vs PC3: 11,680 nM; RWPE2 121,661 nM; PrECs: 37,608 nM; **Fig. S7c**). While serving as an important control, the inactive epimer of LLC0150 (labeled as LLC0150-dead) had no effect on either NSD1/2 protein abundance or the viability of AR-positive prostate cancer cells (**Fig. 4b and Fig. S7d**). LLC0150 also showed marked synergy with enzalutamide—a clinical direct AR antagonist—in killing LNCaP and VCaP cells (**Fig. 4g and Fig. S7e,f**). More impressively, LLC0150 treatment was similarly cytotoxic in cell line models that had acquired resistance to enzalutamide (**Fig. 4h and Fig. S7g**), which is inevitably the case in all enzalutamide-treated patients in the clinic^4,57,58^.

Next, we characterized the cytotoxic effect of LLC0150 in a panel of over 110 human-derived normal and cancer cell lines originating from 22 different lineages (**Table S3**). As expected, hematologic cancers harboring the activating NSD2 E1099K mutation emerged as the most sensitive to treatment with LLC0150 (IC50 ranging from 0.274 - 69.68 nM), which was immediately followed by all the tested AR-positive prostate cancer cell lines (shown in red, **Fig. 4i**). Notably, AR+ disease models showed preferential cytotoxicity to NSD1/2 co-degradation relative to both PC3 and DU145 (AR-negative prostate cancer models) as well as a host of normal human cell lines. Several models of AR+ apocrine breast cancer were also sensitive to LLC0150. In sum, this data suggests that combined loss of the NSD1 and NSD2 paralogs leads to a dramatic, almost complete, loss of the H3K36me2 histone mark and disruption of the AR/FOXA1 neo-enhancer circuitry, resulting in apoptotic cell death of enhancer-addicted prostate cancer cells. This positions NSD1/2 compensatory paralogs as a digenic dependency in AR-driven prostate cancer that can be therapeutically targeted in the advanced, therapy-resistance disease.

## Discussion

In the current clinical regimen of prostate cancer treatment, following surgical resection or radiation, all targeted therapies inhibit the androgen/AR signaling axis^4^. However, how the pro-differentiation AR pathway in normal physiology gets reprogrammed to serve as the central oncogenic node in prostate cancer remains largely unknown. Seminal studies uncovered the global AR chromatin binding profiles to be markedly different between the normal and transformed prostate epithelia^5,6,13–15,59^, and implicated FOXA1 and HOXB13 in mediating AR’s reprogramming^5,14^. However, both FOXA1 and HOXB13 are expressed in the normal prostate epithelial cells, raising the possibility for additional cofactors to underlie the recruitment of AR to cancer-specific enhancer elements. Here, using a functional CRISPR screen, we identify NSD2 as a novel subunit of the AR/FOXA1 enhanceosome. NSD2 is exclusively expressed in the malignant prostate luminal epithelial cells, wherein it enables functional binding of AR at non-canonical, chimeric AR-half motifs, which majorly comprise the AR neo-enhancer circuitries. Consequently, functional inactivation of NSD2 abolishes hallmark cancer phenotypes, while its re-expression in deficient cells restores defining neoplastic features. This positions NSD2 as the first reported *neo*-cofactor of AR that assists oncogenic transcription factors, like FOXA1, HOXB13, and ETS, in redistributing AR on chromatin, thereby unlocking its oncogenic gene programs.

Intriguingly, in motif analyses of the cancer-specific AR cistromes, we also found a modest, yet significant, depletion of the canonical palindromic ARE elements. Most notably, despite a magnitude-fold increase in AR abundance in mCRPC, its loading at *cis*-regulatory elements comprising just the full ARE was largely dismantled. The ARE-containing AR sites were particularly inactivated in mCRPC specimens, as assessed via a dramatic loss of the H3K27Ac mark. This raises an intriguing possibility for the AR transcriptional activity stemming from a subset of canonical elements to rather impede tumor formation and/or progression, which is also consistent with the physiological role of AR as a pro-differentiation factor. In fact, hyper-stimulation of AR activity has anti-proliferative effects in prostate cancer cells^60^, and bipolar androgen therapy involving cyclical inhibition and hyperactivation of AR is being currently tested in advanced patients^61,62^. These are exciting areas for further research.

We further found the loss of NSD2 in prostate cancer cells to lead to a dramatic up-regulation of NSD1 levels, and co-inactivation of both paralogs to be acutely cytotoxic. We uncovered NSD1 and NSD2 to have a compensatory relationship in prostate cancer cells and show that, through disparate mechanisms, they converge on wiring and maintaining the hyper-transcribed oncogenic AR gene programs. While NSD2 directly binds to AR and stabilizes the AR enhanceosome at *de novo* neo-enhancer elements, NSD1 functions as the primary workhorse enzyme for depositing the H3K36me2 mark that antagonizes the PRC2/EZH2 repressive complex. The catalytic activity of PRC2/EZH2 in mCRPC has been shown to repress AR activity^36,37^. Thus, the loss of NSD2 potentially creates an increased dependency on NSD1 in AR-addicted prostate cancer cells, positioning the NSD1/2 paralogs as a newly identified co-vulnerability that can be therapeutically targeted in advanced disease.

We synthesized and characterized a potent dual PROTAC of NSD1 and NSD2 that confirmed co-degradation of these proteins to result in apoptotic cell death in AR-positive prostate cancer. Notably, both NSD1 and NSD2 are recurrently altered in hematological malignancies. NUP98-NSD1 activating fusions are recurrently found in childhood acute myeloid leukemias^63^, while NSD2 translocations and/or catalytic-domain hotspot mutations are detected in multiple myeloma^26–28^ and pediatric acute lymphoblastic leukemia^29–31^, respectively. We found LLC0150 to have the highest potency in NSD2-altered cancers, thus highlighting the potential application of our PROTAC compound in the study of these tumors. Unfortunately, while the LLC0150 PROTAC is a valuable tool to probe NSD1/2 biology *in vitro*, we found that the compound has poor bioavailability *in vivo*. This limited our ability to carry out pre-clinical assessments in animal models in this study. Nonetheless, in our genetic model in which NSD2-null 22RV1 prostate cancer cells stably express the exogenous dTAG-version of NSD2 (**Fig. 3e and S4i**), dTAGV-1-induced degradation of the NSD2 fusion protein led to marked growth inhibition of 22RV1 xenografts, suggesting the potential efficacy of NSD2-targeting agents *in vivo*.

In summary, we identify and functionally characterize NSD2 as an essential cofactor of the AR neo-enhanceosome that is exclusively expressed in prostate cancer cells. NSD2 directly binds to AR and enables its loading at *cis*-regulatory elements harboring chimeric AR-half motifs that constitute over 65% of the malignant AR cistrome. We coalesce these mechanistic insights to propose that AR has two distinct modes of interacting with chromatin: 1) NSD2-independent binding at *cis*-elements harboring canonical, palindromic AREs that are predominantly found in the physiological enhancer circuitry, and 2) NSD2-dependent binding at *cis*-regulatory elements harboring chimeric motifs made of the AR-half site juxtaposed to either FOXA1 or other driver cofactor DNA sequence that distinctively constitute the cancer-specific enhancer/super-enhancer circuitries of AR (**Fig. 5**). Furthermore, we uncover a compensatory relationship between NSD1 and NSD2, positioning them as a novel digenic dependency in AR-positive prostate cancers, and develop an NSD1/2 PROTAC degrader that shows preferential cytotoxicity in AR-addicted and NSD2-altered human cancers. Our findings warrant a focused development of new NSD-targeting therapeutics and evaluation of their efficacy and safety in preclinical and clinical studies.

**Figure 5:**
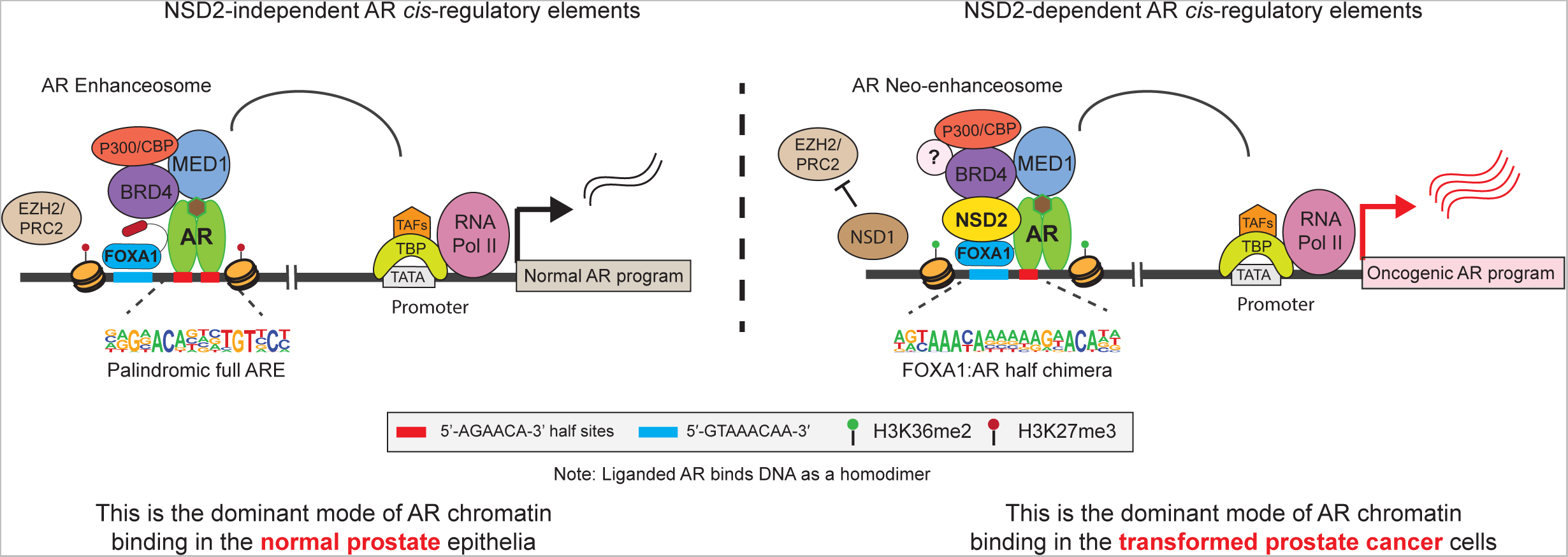
Schema depicting NSD2’s role in loading the AR enhanceosome at tumor-enriched chimeric AR neo-enhancer elements. Chromatin loading of AR in prostate epithelial cells follows two distinct modes of DNA interactions: **Left**, NSD2-independent binding at *cis*-elements harboring the canonical, 15bp palindromic AREs that are predominantly found in the physiological/normal enhancer circuitry, and **Right**: NSD2-dependent loading at *cis*-regulatory elements harboring chimeric AR-half motifs juxtaposed to the FOXA1 sequence that distinctively constitute the cancer-specific enhancer/super-enhancer circuitries.

## Author contributions

A.P., A.M.C., and I.A.A. conceived the study and the experiments; A.P. and S.E. designed and carried out the multi-omics and functional experiments with assistance from J.L., S.E.C., Y.L., L.X., T.H., Y.Q., P.G., M.J., R.R., M.P., E.M., and X.C.; B.K.V. designed and carried out the fragment-based coimmunoprecipitation and functional experiments with assistance from S.A. and R.R; B.K.V. and C.K.D. conducted *in vivo* experiments. A.P. and E.Y. carried out all the bioinformatics analyses with assistance from J.G. and M.A.; Y.Z. analyzed the single-cell RNA-seq datasets; O.T. provided guidance; L.L., C.H., Z.W., and K.D. were involved in the synthesis and structural validation of the LLC0150 compound; A.P. wrote the manuscript and developed the figures with feedback from A.M.C. and I.A.A.; S.E. wrote the Methods section with help from B.K.V., E.Y., Z.W., R.M., and Y.Z. Next-generation sequencing (NGS) related to data shown in Fig. 1c, 3k, 3m, and 4d as well as related supplementary figures was performed at the University of Michigan, while NGS related to data shown in Fig. 1g, 1i, 2a-e, 2i,j, and 4e,f as well as related supplementary figures was carried out at the University of Pennsylvania. All authors discussed the results, provided feedback, and reviewed the final manuscript.

## Acknowledgments

We thank Rui Wang and Fengyun Su from the sequencing team at the Michigan Center for Translational Pathology (MCTP) for generating NGS libraries and coordinating sequencing. We thank Somnath Mahapatra at MCTP for technical assistance in immunofluorescence experiments. We thank Jessica Waninger from MCTP for her technical help with size exclusion chromatography. We also acknowledge critical support from the University of Michigan Biomedical Research Core Facilities, especially the Vector, Flow Cytometry, and Proteomics and Peptide Synthesis Cores. We thank Christopher Vakoc from Cold Spring Harbor Laboratories for generously sharing the domain-focused epigenetics sgRNA library. We thank A. Heller, O. Rivera, A. Pawar, and R. Natesan from the University of Pennsylvania for providing technical and bioinformatics assistance. This research was supported by the following mechanisms: Prostate Cancer Foundation (PCF), Prostate Specialized Programs of Research Excellence (SPORE) Grant P50-CA186786, National Cancer Institute Outstanding Investigator Award R35-CA231996, National Cancer Institute P30-CA046592, and National Cancer Institute R00-CA187664. A.P. is supported by the NIH/NCI K00 fellowship (K00-CA245825), Michigan SPORE Career Enhancement Program, PCF Young Investigator Award, and Rogel Fellowship. L.X. is supported by a Department of Defense Prostate Cancer Research Program Idea Development Award (W81XWH-21-1-0500) and a PCF Young Investigator Award. A.M.C. is a Howard Hughes Medical Institute Investigator, A. Alfred Taubman Scholar, and American Cancer Society Professor. I.A.A. is supported by grants from the National Institute of Health and Department of Defense (R01-CA249210-0 and W81XWH-17-0404).

## Figure Legends

**Figure S1: Generation and characterization of the endogenous mCherry-PSA AR reporter cell lines.**

a. Schematic representation of the workflow of LNCaP-mCherry-PSA AR reporter cell line generation.
b. DNA gel electrophoresis image showing the exogenously inserted mCherry amplicon in the LNCaP-mCherry-PSA lines. Clones 1 and 2 were used for the functional CRISPR screen.
c. Sanger sequencing chromatograms of the PCR amplicon from reporter cells in panel (b) showing the KLK3/PSA gene promoter and exon 1 start codon junctions.
d. Representative brightfield and mCherry immunofluorescence images of the LNCaP-mCherry-PSA clone 1 treated with (top) AR-targeting siRNA or ASO’s (siAR and ASO AR respectively) or enzalutamide (bottom left). Reporter cells were also serum starved for 48h and stimulated with DHT (10nM for 12h) to showcase gain in signal (bottom right).
e. Immunoblots of noted proteins in LNCaP reporter cells as in panel (d).
f. Expression (qPCR) of noted genes in reporter monoclones treated as in panel (d) to manipulated AR signaling.
g. Immunoblots of noted proteins, including the exogenously introduced mCherry protein, in LNCaP reported cells treated with AR-targeting epigenetic drugs. Total H3 is used as a loading control.
h. Next-generation sequencing-based abundance of sgRNAs in the epigenetic-focused library used in the CRISPR screen highlighting some of the known epigenetic regulators of AR.

**Figure S2: NSD2 transcript and protein expression in primary patient specimens.**

a. Immunoblot of labeled proteins in a collection of AR+ and AR- prostate cell lines. GAPDH and H3 are used as a loading control.
b. Quantitative-PCR (qPCR) of KLK3 expression in LNCaP NSD2 WT and KO cells stimulated with R1881 for 12 and 24 hours. HPRT1 is used as a loading control.
c. Immunoblot of labeled proteins in LNCaP NSD2 WT and KO cells stimulated with DHT for 30 hours.
d. UMAP plots from patient-matched normal and primary prostate cancer single-cell RNA-seq data.
e. NSD2 and PCA3 transcript expression in patient-matched normal and primary prostate cancer luminal epithelial cells (pseudo-bulk analyses from single cell data; n=15, Wilcoxon test).
f. Representative immunohistochemistry (IHC) images of NSD2 in normal, primary prostate cancer, and metastatic castration-resistant prostate cancer patient specimens.
g. Boxplot showing RNA expression of labeled genes in primary prostate cancer specimens (TCGA cohort) stratified by the Gleason score. (Kruskal-Wallis test).

**Figure S3: Motif characterization of the NSD2-enabled AR neo-cistrome in prostate cancer cells.**

a. *Left*: Schematic representation of the half-motif enrichment analysis. *Right*: Motif enrichment plot of AR-half motifs with neighboring motifs of other transcription factors at NSD2-dependent and independent AR sites in LNCaP cells.
b. Venn diagram showing overlaps between AR ChIP-seq sites in LHSAR cells with LacZ (control), FOXA1, HOXB13, FOXA1+HOXB13 overexpression.
c. Venn diagram showing overlap of AR cistromes (ChIP-seq) in normal prostate, primary prostate cancer, and castration-resistant prostate cancer specimens. (Pomerantz et. al. 2015 & 2020)
d. Motif fold-change heatmap in normal, primary cancer, and castration-resistant prostate cancer specimens.
e. Fold change and significance of HOMER motifs enriched within mCRPC cancer-specific AR sites over normal tissue-specific AR elements (data from Pomerantz, 2015 and 2020).
f. Boxplot showing H3K27Ac ChIP-seq read density at sites containing the ARE or the FOXA1:AR motif in normal and tumor patient samples (normal prostate, n=7; primary prostate cancer, n=13; castration-resistant prostate cancer - CRPC, n=15). In box plots, the center line shows the median, box edges mark quartiles 1-3, and whiskers span quartiles 1-3 土 1.5 * interquartile range.
g. ChIP-seq read density tracks of AR and H3K27Ac within the SLC45A3 and TMPRSS2 loci in NSD2 WT and NSD2 KO LNCaP cells. Super-enhancer clusters are highlighted in a grey box.

**Figure S4: Molecular characterization of the NSD2-rescued prostate cancer cells.**

a. *Left*: Immunoblots of noted proteins upon long-term treatment with control siRNA (siNC) or NSD2-targeting siRNA (siNSD2). *Right*: Immunoblot of NSD2 in LNCaP and 22RV1 cells treated with a control sgRNA (NSD2 WT) or sgRNA targeting NSD2 (NSD2 KO). GAPDH and total histone H3 are used as loading controls.
b. *Left:* Representative images of colonies of control or NSD2-null LNCaP cells. *Right*: ImageJ-based quantification of stained colonies shown in the images to the left (two-sided t-test).
c. *Top*: Tumor volumes of 22RV1 parental or NSD2-KO+HA-tagged NSD2-L cell line-derived xenograft in mice. *Bottom*: Immunoblot of noted proteins from the 22RV1 xenograft tumors from the growth experiment at endpoint.
d. *Top, left*: Representative images of Boyden chambers showing invaded cells stained with calcein AM dye in the LNCaP NSD2 WT and KO or NSD2-L rescued lines. *Right*: Quantified fluorescence signal from invaded cells (two-way ANOVA and Tukey’s test) *Bottom*: Immunoblot of listed proteins in wildtype or NSD2 KO LNCaP cells with stable exogenous overexpression of NSD2-L and/or NSD2-S isoforms. eGFP is used as control.
e. Immunoblots showing expression of the KLK3 (aka PSA) protein in the eGFP or NSD2 overexpressing LNCaP cells.
f. Immunoblots of noted proteins in the NSD2 wildtype (WT) or NSD2-knocked out (KO) LNCaP cells that are rescued with exogenous WT or SET domain-deleted NSD2 mutants.
g. Immunoblots of noted proteins in LNCaP cells overexpressing the hyper-catalytic NSD2 SET domain E1099K mutant.
h. Immunoblots of noted proteins in the 22RV1 NSD2-FKBP12-F36V engineered cell lines upon dTAG-13 treatment (0.5uM for 6h).
i. *Left*: Box plot showing tumor weights of 22RV1+NSD2-FKBP12-F36V xenografts at endpoint (day 18) upon treatment with dTAGv-1. *Right*: Tumor images at the endpoint from the animal growth studies.
j. Heatmap showing expression (qRT-PCR) of AR target genes in the 22RV1 NSD2-KO + NSD2-FKBP12-F36V cell line after 12h or 24h of treatment with dTAG-13.
k. Immunoblots of noted proteins in size-exclusion chromatography fractions of nuclear lysate extracted from wild-type LNCaP cells. Fractions containing the AR protein are marked.

**Figure S5: Fragment-based NSD2–AR co-immunoprecipitation and characterization of the NSD paralog knockout prostate cancer cells.**

a. Immunoblots of indicated proteins upon co-immunoprecipitation of AR in prostate cancer cells.
b. Immunoblots of indicated proteins upon immunoprecipitation of exogenously expressed Halo-tagged full-length AR protein in HEK293FT cells that express HA-tagged NSD2 fragments. Both input (left) and immunoprecipitation (right) blots are shown.
c. *Left*: Immunoblots of HA-tag-based immunoprecipitation of full length NSD2 in HEK293FT cells that express the Halo-tagged AR protein fragments. *Right*: Immunoblots of Halo-tag-based immunoprecipitation of the DNA-bidning domain (DBD) of AR in HEK293FT cells that overexpress different HA-tagged NSD2 fragments. For both experiments, input and immunoprecipitation blots are shown.
d. Heatmap of AR upregulated genes (z-score) in NSD1 or NSD3 knockout (KO) LNCaP cells.
e. GSEA plots for AR-regulated genes using the fold change rank-ordered genes from LNCaP NSD3 knock-out (NSD3 KO) vs control cell lines. DEGS, differentially expressed genes.
f. Immunoblot of indicated proteins in NSD1 or NSD3-deficient LNCaP cells.
g. GSEA net enrichment score (NES) plot of downregulated hallmark pathways in LNCaP NSD1 knocked out (KO) vs wild-type control cells.
h. Immunoblot of indicated proteins upon treatment with NSD1 and NSD2-targeting siRNAs (labelled as siNSD1 and siNSD2) independently or in combination in VCaP cells.
i. Dependency map (DepMap) plots showing the dependency scores for NSD1, NSD2, NSD3, and POLD2 (positive control; pan-essential gene) across cell lines from distinct originating tissues. The red dotted line indicates pan-essentiality z-score cutoff.

**Figure S6: Mechanistic characterization of the NSD1/2 PROTAC degrader LLC0150.**

a. Immunoblots of indicated proteins in VCaP and LNCaP cells pre-treated with bortezomib, thalidomide, or VL-285 followed by treatment with LLC0150 at noted concentrations.
b. Heatmap of relative abundance of several PWWP-domain-containing proteins detected via Tandem Mass Tag (TMT) based quantitative MS upon 12h treatment with LLC0150 in VCaP cells.
c. Genome-wide changes in FOXA1 ChIP-seq peaks in LNCaP cells treated with LLC0150 (2uM for 48h).
d. Immunoblots of noted proteins in whole-cell or chromatin lysates from VCaP and LNCaP cells treated with LLC0150 (2uM) for 24h.
e. Heatmap of z-score normalized expression (qRT-PCR) of AR target genes in LNCaP and VCaP cells treated with LLC0150 followed by DHT stimulation (10nM for 24h). Treatment with DHT alone is used a control.
f. Read density ChIP-seq tracks of AR, FOXA1, and H3K27Ac within the TMPRSS2 super-enhancer in LNCaP cells treated with LLC0150 (2uM for 24h). Super-enhancer cluster is highlighted in a grey box.

**Figure S7: Transcriptomic effect and drug synergism of LLC0150 in prostate cancer cells.**

a. GSEA plots for E2F, G2M, and apoptosis pathway genes using the fold change rank-ordered genes from the LLC0150 vs DMSO treated LNCaP (left) or VCaP (right) cell lines. DEGS, differentially expressed genes.
b. Immunoblot of noted proteins in LNCaP cells treated with LLC0150 (2uM for 72h), dead-analog (LLC0150-dead), or the warhead alone (UNC6934). LLC0149 is an independent NSD1/2 PROTAC.
c. Dose-response curves of LLC0150 in normal prostate, AR-positive, or AR-negative prostate cancer cell lines at the indicated concentrations for five days.
d. Dose-response curves of LLC0150 and its inactive epimer control (LLC0150-dead) in LAPC4 and VCaP cell lines.
e. Percent growth inhibition (Cell-titer Glo) of VCaP cells upon co-treatment with varying concentrations of LLC0150 and enzalutamide for 5 days.
f. 3D synergy plots of LLC0150 and enzalutamide co-treated LNCaP and VCaP cells. Red peaks in the 3D plots denote synergy with the average synergy scores noted above.
g. Dose-response curves of LLC0150 in LNCaP parental and enzalutamide-resistant cell lines at varying concentrations for five days. Half maximal inhibitory concentration (IC50) at noted.

**Table S1:** List of known motifs enriched in the NSD2-dependent AR elements along with their HOMER statistics.

**Table S2:** List of known motifs enriched in the NSD2-independent AR elements along with their HOMER statistics.

**Table S3:** Antiproliferative half-maximal inhibitory concentration of LLC0150 across human-derived normal and cancer cell lines.

## Materials and Methods

### Cell lines

Most cell lines were purchased from the American Type Culture Collection (ATCC) and were cultured following ATCC protocols. For all experiments, LNCaP and 22RV1 cells were grown in RPMI 1640 medium (Gibco) and VCaP cells in DMEM with Glutamax (Gibco) medium supplemented with 10% full bovine serum (FBS; Invitrogen). HEK293 cells were grown in DMEM (Gibco) medium with 10% FBS. All cells were grown in a humidified 5% CO2 incubator at 37 °C. Mycoplasma and cell line genotyping were performed once a fortnight and every month respectively at the University of Michigan Sequencing Core using Profiler Plus (Applied Biosystems). Results from these were compared with corresponding short tandem repeat profiles in the ATCC database to authenticate their identity.

### Antibodies

For immunoblotting, the following antibodies were used: NSD1 (NeuroMab: 75-280); NSD2 (Abcam:ab75359); NSD3 (Cell Signaling Technologies: 92056S); KLK3/PSA (Dako:A0562); FKBP5(Cell Signaling Technologies: 12210); NKX3-1 (Cell Signaling Technologies:83700S); FOXA1 N-terminal (Cell Signaling Technologies: 58613S; Sigma-Aldrich: SAB2100835); FOXA1 C-terminal (Thermo Fisher Scientific: PA5-27157); AR (Millipore: 06-680); AR (Abcam: ab133273); H3 (Cell Signaling Technologies: 3638S); GAPDH (Cell Signaling Technologies: 3683); H3K27me3(Millipore: 07-449); H3K36me2 (Cell Signaling Technologies: 2901S); H3K27Ac (Active Motif, Cat#39336); Phospho-AR (Ser-81) (Millipore, Cat# 07-1375-EMD); HALO (Thermo Scientific, Cat# G9281); HA (Cell Signaling Technologies, Cat# 3724S); His (Cell Signaling Technologies, Cat#2365). ChIP-seq assays were performed using the following antibodies: FOXA1 (Thermo Fisher Scientific: PA5-27157); AR (Abcam: ab133273); and H3K27Ac (Active Motif, Cat#39336).

### Western blot and immunoprecipitation

Cells were lysed in RIPA buffer (ThermoFisher Scientific) supplemented with cOmpleteTM protease inhibitor cocktail tablets (Sigma-Aldrich). Protein concentrations were measured using Pierce BCA Protein Assay Kit (ThermoFisher Scientific). For immunoblotting, equal amounts of protein were resolved in either a NuPAGE 3 to 8%, Tris-Acetate Protein Gel (ThermoFisher Scientific) or NuPAGE 4 to 12%, Bis-Tris Protein Gel (ThermoFisher Scientific) and incubated overnight with primary antibodies. Following incubation with HRP-conjugated secondary antibodies (BioRad), membranes were imaged on an Odyssey CLx Imager (LiCOR Biosciences).

For immunoprecipitation (IP) experiments, nuclear or whole cell protein extracts were obtained from cells using NE-PER nuclear extraction kit (Thermo Scientific, 78835) or RIPA buffer, respectively. The nuclear pellet was then lysed in an IP buffer by sonication. Nuclear lysates (0.5-2.0 mg) were pre-cleared by incubation with protein G Dynabeads (Life Technologies, 10003D or Thermo Scientific, 14321D) for 1 h on a rotator at 4°C. Next, an antibody (2-5 μg) was added to the pre-cleared lysates and incubated on a rotator at 4°C overnight. The following day, Protein G Dynabeads were added to the nuclear lysate-antibody mix for 1 h. Beads were washed twice in IP or RIPA buffer containing 300 mM NaCl, resuspended in 40 μL of 2x loading buffer, boiled at 100°C for 5 min, and subjected to SDS-PAGE and immunoblotting. For Halo-tag immunoprecipitation, the cells were lysed in NP-40 buffer (0.1% NP-40, 15 mM Tris–HCl pH7.4, 1 mM EDTA, 150 mM NaCl, 1 mM MgCl2, 10% Glycerol) containing protease and phosphatase inhibitors and the lysates were incubated with anti-Halo M2 beads for 2 h at 4°C. After that, SDS–PAGE, and immunoblotting was performed as described above. Densitometric quantification was performed using the Image J software and Image Lab Software.

All co-immunoprecipitation experiments were performed at least twice. Statistical significance of immunoblotting data was assessed using a one-way analysis of variance (ANOVA) and a Tukey’s honestly significantly different (HSD) post hoc test for multiple comparisons, or unpaired Students t-test for pairwise comparisons. In all cases, P values of <0.05 were considered statistically significant and are indicated with *P < 0.05, **P < 0.01, ***P < 0.005, ****P < 0.001. Statistical analysis was performed in GraphPad Prism.

### Cell free protein-protein interaction studies

In vitro protein expression was carried out by cloning the desired expression cassettes downstream of a Halo- or His-tag to produce fusion proteins. Briefly, AR-DBD was subcloned in pFN21K containing Halo tag and NSD2-HMGa was cloned in pcDNA4c containing His tag. After cloning, the fusion proteins were expressed using the cell-free transcription and translation system (Cat. # L4140, Promega) following the manufacturer’s protocol. For each reaction, protein expression was confirmed by Western blot.

A total of 10μl cell-free reaction containing halo- and His-tag fusion proteins were incubated in PBST (0.1% tween) at 4°C overnight. Ten microliter HaloLink beads (Cat. #G931, Promega) were blocked in BSA at 4°C for overnight. After washes with PBS, the beads were mixed with AR-NSD2-HMGa and TM mixture and incubated at RT for 1hr. Halolink beads were then washed with PBST for 4 times and eluted in SDS loading buffer. Proteins were separated on SDS gel and blotted with anti-His Ab (CST).

### Generation of the endogenous AR reporter and functional CRISPR screen

LNCaP cells were treated with CRISPR guides targeting the promoter-5’UTR junction of KLK3/PSA and a knock-in repair template containing the mCherry coding sequence, P2A cleavage site (as represented in Fig. 1 and Fig. S1) along with 750bp of flanking KLK3 homologous sequences were transfected at a ratio of 1:3 (guide: repair-template) using Lipofectamine 3000 reagent (Thermo Fisher Scientific, Cat# L3000001). To ensure preferential homologous recombination, non-homologous end joining repair (NEHJ) was inhibited using SCR7 (2uM for 24 hours). Post transfection, cells were subject to puromycin selection for three days, and surviving mCherry positive cells were FACS sorted to establish monoclonal reporter lines. The top 75% of mCherry positive cells were sorted to ensure robust and uniform mCherry expression. All intended repair junctions were amplified and Sanger sequenced to confirm sequence integration.

KLK3/PSA gRNA sequence used is as follows: 5’-CACCGAAGACAACCGGGACCCACA-3’

Two monoclones, hereafter named as Clone 1 and 2, were used for the downstream CRISPR screen with stable viral integration of the active Cas9 enzyme. Briefly, the cells were treated with an epigenetic guide library (a kind gift from Dr. Christopher Vakoc), puromycin selected, and incubated for eight days, after which they were stimulated with 1nM DHT for 16 hours and sorted into mCherry high and low populations. Input fractions were also collected at Day 3 post-puromycin selection for guide enrichment analysis. Genomic DNA from input and sorted cells was extracted (DNeasy Blood and Tissue kit) and integrated guide RNA sequences were amplified using common primers, with the resulting amplicon pool submitted for next-generation sequencing.

### Colony formation assays

For the colony formation assay, approximately 10,000 cells/well in six-well plates (n=3) were seeded and treated with the required drugs/compounds or vehicle for 12-14 days. Media was replenished every 3-4 days. Colonies were fixed and stained using 0.5% (w/v) crystal violet (Sigma, C0775) in 20% (v/v) methanol for 30 min, washed with distilled deionized water, and air-dried. After scanning the plate, the stained wells were destained with 500 μL 10% acetic acid, and the absorbance was determined at 590 nm using a spectrophotometer (Synergy HT, BioTek Instruments, Vermont-USA).

### Cellular protein fractionation assays

Chromatin-bound proteins were extracted following a protocol previously described (Ur Rasool et al., 2019). In brief, 10 million cells were collected, washed with DPBS, and resuspended in 250 μl Buffer A (10 mM HEPES pH 7.9, 10 mM KCl, 1.5 mM MgCl_2_, 0.34 M sucrose, 10% glycerol, 1 mM DTT) supplemented with 0.1% TritonX-100. After incubation on ice for 10 min, the nuclear pellet was collected by centrifugation at 1300 x g for 5 min at 4°C, washed in Buffer A, and resuspended in Buffer B (3 mM EDTA, 0.2 mM EGTA, 1 mM DTT) with the same centrifugation settings, and incubated on ice for 30 min. The chromatin pellet was collected by centrifugation at 1700 x g for 5 min at 4°C, washed and resuspended in Buffer B with 150 mM NaCl, and incubated on ice for 20 min. After centrifugation at 1700 x g for 5 min to remove proteins soluble in 150 mM salt concentrations, the pellet was then incubated in Buffer B with 300 mM NaCl on ice for 20 min and centrifuged again at 1700 x g to obtain the final chromatin pellet. The chromatin pellet was dissolved in a sample buffer, sonicated for 15 seconds, and boiled at 95°C for 10 min. Immunoblot analysis was conducted on samples as described previously. All buffers were supplemented with Pierce protease inhibitor and Halt protease & phosphatase inhibitors.

### RNA isolation and quantitative real-time PCR

Standard protocols from the miRNeasy Mini kit (Qiagen) was used to extract total RNA with the inclusion of on-column genomic DNA digestion step using the RNase-free DNase Kit (Qaigen). RNA concentration was estimated using the NanoDrop 2000 spectrophotometer (ThermoFisher Scientific), and 1ug of total RNA was used for complementary DNA (cDNA) synthesis using the SuperScript III Reverse Transcriptase enzyme (ThermoFisher Scientific) following manufacturer’s instructions. 20ng of cDNA was used for each polymerase chain reaction (PCR) using the FAST SYBR Green Universal Master Mix (ThermoFisher Scientific), and every sample was quantified in triplicates. Gene expression was normalized and calculated relative to GAPDH and HPRT1 (loading control) using the delta-delta Ct method and normalized to the control group for graphing. Quantitative PCR (qPCR) primers were designed using the Primer3Plus tool (http://www.bioinformatics.nl/cgi-bin/primer3plus/primer3plus.cgi) and synthesized by Integrated DNA Technologies.

Primers used in this study are listed below:

*GAPDH:* forward (F), TGCACCACCAACTGCTTAGC and reverse (R), GGCATGGACTGTGGTCATGAG; *FKBP5*: F, TCTCATGTCTCCCCAGTTCC and R,TTCTGGCTTTCACGTCTGTG; *TMPRSS2:* F,CAGGAGTGTACGGGAATGTGATGGT and R,GATTAGCCGTCTGCCCTCATTTGT; *AR:* F,CAGTGGATGGGCTGAAAAAT and R,GGAGCTTGGTGAGCTGGTAG; *NKX3-1:* F,ACGTCCTTCCTCATCCAGGACA and R,AGGGCGCCTGAAGTGTTTTC; SLC45A3: F, TCGTGGGCGAGGGGCTGTA and R,CATCCGAACGCCTTCATCATAGTGT; ZBTB16: F, CAGTTTTCGAAGGAGGATGC and R,CCCACACAGCAGACAGAAGA; KLK3: F, ACGCT GACAGGGGGCAAAAG and R, GGGCAGGGCACA TGGTTCACT

### siRNA-mediated gene knockdown

Cells were seeded in a 6-well plate at the density of 100,000–250,000 cells per well. After 12 hours, cells were transfected with 25nM of gene-targeting ON-TARGETplus SMARTpool siRNAs or non-targeting pool siRNAs as negative control (Dharmacon) using the RNAiMAX reagent (Life Technologies; Cat#: 13778075) on two consecutive days, following manufacturer’s instructions. Both total RNA and protein was extracted on day 3 (total 72h) to confirm efficient (>80%) knockdown of the target genes. For the siRNA-treated VCaP DMSO/EPZ-6438 RNA-seq experiment (Fig. 3k), cells were pre-treated with control siRNA (siNC) or siRNA targeting NSD1, NSD2 or NSD1/2 (siNSD1, siNSD2) for 30 days, followed by 72hours of EPZ-6438 treatment.

Catalog numbers and guide sequences (5’ to 3’) of siRNA SMARTpools (Dharmacon) used are: non-targeting control (cat. no. D-001810-10-05; UGGUUUACAUGUCGACUAA, UGGUUUACAUGUUGUGUGA,UGGUUUACAUGUUUUCUGA,UGGUUUACAUGUUUUCCUA); NSD1:(Cat#L-007048-00-0005:GGACGAGAAUUCUUUGAUU,GAACAGAAGUAGUACCAAU,GAUCAAAGCCUUCAUCCAA,GC CGAGAGCUGUUGAGAAA);

NSD2:(Cat#L-006571-01-0005:GGUCCAAAGUGUCGGGUUA,UGUCAGUGGAGGAGCGGAA,GGAGCAGGGCCUUGUCGAA, CCGGGUGUUUAAUGGAGAA);

NSD3(Cat#L-012875-00-0005: GGUUGACACUGUAUCAGAA, GAACGUGCUCAGUGGGAUA, GCUUGAGGUUCAUACUAAA, GUCCACUGGUGUUAAGUUU)

### CRISPR-Cas9-mediated gene knockout

To CRISPR knockout genes, cells were seeded in a 6-well plate at the density of 200,000–300,000 cells per well and infected with viral particles carrying the lentiCRISPR-V2 constructs coding either non-targeting control guides (sgNC) or single guide RNAs (sgRNA)-targeting NSD1 and NSD2. This was followed by 3 days of puromycin selection, after which proliferation assays were carried out as described below. The lentiCRISPR-V2 vector was a gift from Dr. Feng Zhang’s lab (Addgene plasmid # 52961).

sgRNA sequences used are as follows: sgNC#1:5′-GTAGCGAACGTGTCCGGCGT-3′; sgNC#2:5′-GACCGGAACGATCTCGCGTA-3′; sgNSD2-SET:5’CACCGAAAGTCCAGATCTACACAG3’; sgNSD2-Long:5’CACCGCCCCGTTGCCTATCACAGCG3’; sgNSD1#1:5’CACCGAGTGGCGTAGACTTTTTCTT3’; sgNSD1#2:5’CACCGCTATCGGCAGTACTACGTGG3’; sgNSD1#3:5’CACCGTATGTGAGTTACTAATGAG3’; sgNSD1#4:5’ CACCGTAAGAGAGTTCCTTCAACT 3’; sgNSD3#1: 5’ CACCGGATACCCATTTGTGAGTGG 3’; sgNSD3#2: 5’ CACCGTTTGTGGTACCGAAGGAGG 3’

### Proliferation assays

For siRNA growth assays, cells were directly plated in a 96-well plate at the density of 2,500–8,000 cells per well and transfected with gene-specific or non-targeting siRNAs, as described above, on day 0 and day 1. Every treatment was carried out in six independent replicates. CellTiter-Glo reagent (Promega) was used to assess cell viability at multiple time points after transfection, following the manufacturer’s protocol. Data were normalized to readings from siNC treatment on day 0 and plotted as relative cell viability to generate growth curves. Alternatively, for CRISPR sgRNA growth assays, CRISPR-edited cells were seeded into a 96-well plate at 1,000 - 5,000 cells per well with six replicates per group.

### Matrigel invasion assay

LNCaP CRISPR clones were grown in 10% CSS-supplemented medium for 48 h for androgen starvation. A Matrigel-coated invasion chamber was used, which was additionally coated with a light-tight polyethylene terephthalate membrane to allow for fluorescent quantification of the invaded cells (Biocoat: 24-well format, no. 354166). Fifty thousand starved cells were resuspended in serum-free medium and were added to each invasion chamber. Twenty percent FBS-supplemented medium was added to the bottom wells to serve as a chemoattractant. After 12 h, medium from the bottom well was aspirated and replaced with 2 μg/ml Calcein-green AM dye (Thermo Fisher Scientific; C3100MP) in 1× HBSS (Gibco) and incubated for 30 min at 37 °C. Invasion chambers were then placed in a fluorescent plate reader (Tecan-Infinite M1000 PRO), and fluorescent signals from the invaded cells at the bottom were averaged across 16 distinct regions per chamber to determine the extent of invasion. For rescue experiments, stable lines overexpressing the NSD2 isoforms were generated. Briefly, to LNCaP NSD2 KO lines, GFP or NSD2-Long isoform containing viruses were added. These lines were then used to perform the invasion assay as described above.

### Size exclusion chromatography

Nuclear lysate from VCaP or LNCaP cells were extracted using the NE-PER nuclear extraction kit (Thermo Scientific) and dialyzed against the FPLC buffer (20 mM Tris-HCl, 0.2 mM EDTA, 5mM MgCl2, 0.1 M KCl, 10% (v/v) glycerol, 0.5 mM DTT, 1 mM benzamidine, 0.2 m MPMSF, pH7.9). 5 mg of nuclear protein was concentrated in 500ul using a Microcon centrifugal filter (Millipore) and then applied to a Superose 6 size exclusion column (10/300 GL GE Healthcare) pre-calibrated using the Gel Filtration HMW Calibration Kit (GE Healthcare). 500 μl elute was collected for each fraction at a flow rate of 0.5ml/min, and eluted fractions were subjected to SDS-PAGE and western blotting.

### RNA-seq and analysis

RiboErase RNA-seq libraries were prepared using 200–1,000 ng of total RNA. Ribosomal RNA was removed by enzymatic digestion of the specific probe-bound duplex rRNA (KAPA RNA Hyper+RiboErase HMR, Roche) and then fragmented to around 200-300bp with heat in the fragmentation buffer. Following this, double-stranded cDNA was generated, and end-repair and ligation was performed using New England Biolabs (NEB) adapters. Final library preparation was performed by amplification with the 2x KAPA HiFi HotStart mix and NEB dual barcode following the manufacturer’s protocol. Library quality was measured on an Agilent 2100 Bioanalyzer for product size and concentration. Paired-end libraries were sequenced with the Illumina HiSeq 2500, (2 × 100 nucleotide read length) with sequence coverage to 15–20M paired reads.

RNA data was first processed using kallisto (version 0.46.1)^64^. Then analysis was performed in R, first read counts were normalized and filtered (counts >10) using EdgeR^65^ (edgeR_3.39.6), and differential expression was performed using Limma-Voom (limma_3.53.10)^66^. Gene Set Enrichment Analysis (GSEA) was performed using fgsea (fgsea_1.24.0)^67^ and comparisons were made to several signatures, including an experimentally derived AR signature, the human hallmark MsigDB signatures (www.gsea-msigdb.org), and the hallmark androgen response signature (HALLMARK_ANDROGEN_RESPONSE.v7.5.1.gmt). In addition, R packages tidyverse, gtable, gplots, ggplot2, and EnhancedVolcano (EnhancedVolcano_1.15.0) were also used for generating summary figures (R version 4.2.1 ^68–70^).

### ChIP–seq and data analysis

Chromatin immunoprecipitation experiments were carried out using the Ideal ChIP-seq Kit for Transcription Factors or Histones (Diagenode) as per the manufacturer’s protocol. Chromatin from 2 × 10^6^ cells (for transcription factors) and 1×10^6^ cells (for histones) was used for each ChIP reaction with 4 or 2 μg of the target protein antibody, respectively. In brief, cells were trypsinized and washed twice with 1× PBS, followed by cross-linking for 8 min in 1% formaldehyde solution. Crosslinking was terminated by the addition of 1/10 volume 1.25 M glycine for 5 min at room temperature followed by cell lysis and sonication (Bioruptor, Diagenode), resulting in an average chromatin fragment size of 200 bp. Fragmented chromatin was then used for immunoprecipitation using various antibodies, with overnight incubation at 4 °C. ChIP DNA was de-crosslinked and purified using the iPure Kit V2 (Diagenode) using the standard protocol. Purified DNA was then prepared for sequencing as per the manufacturer’s instructions (Illumina). ChIP samples (1–10 ng) were converted to blunt-ended fragments using T4 DNA polymerase, *Escherichia coli* DNA polymerase I large fragment (Klenow polymerase), and T4 polynucleotide kinase (New England BioLabs (NEB)). A single adenine base was added to fragment ends by Klenow fragment (3′ to 5′ exo minus; NEB), followed by ligation of Illumina adaptors (Quick ligase, NEB). The adaptor-ligated DNA fragments were enriched by PCR using the Illumina Barcode primers and Phusion DNA polymerase (NEB). PCR products were size-selected using 3% NuSieve agarose gels (Lonza) followed by gel extraction using QIAEX II reagents (Qiagen). Libraries were quantified and quality checked using the Bioanalyzer 2100 (Agilent) and sequenced on the Illumina HiSeq 2500 Sequencer (125-nucleotide read length).

ChIP-seq analysis was carried out by first assessing reads and performing trimming using Trimmomatic version 0.39 (settings TruSeq3-PE-2.fa:2:30:10, minlen 50)^71^. Paired end reads were aligned to hg38 (GRCh38) human genome reference using bwa (“bwa mem” command with options -5SP -T0, version 0.7.17-r1198-dirty) ^72^. Alignments were then filtered using both samtools^73^ (quality score cutoff of 20) and picard ^74^ MarkDuplicates (removed duplicates). Peak calling was performed using MACS2 ^75^ using narrowpeak setting for narrow peaks and a second set for broad peaks (e.g., H3K27Ac, --broad -B -- cutoff-analysis --broad-cutoff 0.05 --max-gap 500). Finally, bedtools^76^ was used to remove blacklisted regions of the genome from the peak list (Encode’s exclusion list ENCFF356LFX.bed). UCSC’s tool wigtoBigwig was used for conversion to bigwig formats^77^.

The castration-resistant prostate cancer (CRPC), normal, and primary prostate cancer (PCa) dataset fastqs were pulled from GEO from Baca et al and Pomerantz et al^5,78^ (GSE130408 and GSE70079) and processed using our previously described ChIP-seq pipelines, adapted for single end reads. Picard utility MergeSamFiles was utilized to combine the aligned bam files. Samtools view -bs was used to subsample the combined bam file to a depth of approximately 100M reads for AR datasets and 250M for H3K27Ac datasets. Peak calling was then repeated using MACS2 callpeak, and output bedgraphs were converted using wigToBigWig. Motif enrichment ratios between primary PCa (or CRPC) to normal prostate tissues were calculated and plotted in Fig. 2g and Fig. S3e. Normalized read density at specific motifs is plotted in Fig. 2h and Fig. S3f.

### Overlap analysis in ChIPseq experiments

Peak lists from MACS were compared between samples using R package ChIPpeakAnno ^79–81^. Peaks within 500bp of each other were reduced to single peaks. Overlaps were calculated using settings maxgap=-1L, minoverlap=0L, ignore.strand=TRUE, connectedPeaks=c(‘keepAll’, ‘min’, ‘merge’). Comparisons of enrichment sites to the known gene database (TxDb.Hsapiens.UCSC.hg38.knownGene) were performed using R package ChIPseeker. A distance of +/− 1kb was used to assess relative distance from gene regions.

### HOMER motif calling

*De novo* and known motif enrichment analysis was performed using HOMER ^47,82^. Custom motif matrices were generated manually, then assigned score thresholds using HOMER’s utility seq2profile, allowing for two mismatches. This setting was chosen after iteratively comparing performance with the pre-existing FOXA1:AR motif. Further customization was achieved by checking for presence of motif elements with different spacings, ranging from 0-8 ‘N’s added between elements, and flipping the order of elements in each of these: FOXA1-ARE, ARE-FOXA1, FOXA-N-ARE, ARE-N-FOXA1, FOXA1-NN-ARE, ARE-NN-FOXA1, etc.

Custom motifs were then further validated using XSTREME ^83^ from the MEME Suite ^82^ to check for additional configurations and variations in padding between motif elements.

### Enrichment heatmaps

The software Deeptools was used to generate enrichment plots and read density heatmaps. A reference point parameter of +/− 2.5kb for histone signals and +/− 1.5kb for AR/FOX signals was used. Other settings included using ‘skipzeros’, ‘averagetype mean,’ and ‘plotype se’. The Encode blacklist ENCFF356LFX was used ^84^.

### Motif and signal plots

Sushi (Sushi_1.32.0) package in R was used to layer signal tracks. The plotBedgraph(), plotGenes(), plotBed() functions were used with output from ChIP-Seq alignments and output from HOMER motif enrichment analysis^85^.

### Super-enhancer analysis

Super-enhancer regions were identified with findPeaks function from HOMER (version v.4.10) ^47^ using options “-style super -o auto”. In addition, the option “-superSlope −1000” was added to include all potential peaks, which were used to generate the super-enhancer plot (super-enhancer score versus ranked peaks). The slope value of greater than or equal to 1 was used to identify super-enhancer clusters. The input files to findPeaks were tag directories generated from alignment files in SAM format with makeTagDirectory function from HOMER. Super-enhancer scores were plotted using the normalized tag count values between the datasets.

### DepMap analysis

To perform the DepMap analysis, we used a BioConductor Cancer Dependency map R data package. This package can directly access the Broad Institute’s CRISPR Achilles DepMap portal. A detailed guide and code to using this package is provided in the reference paper. Briefly, we inputted the CRISPR data using the EH3797 experimental hub. Utilizing the dependency score option, we plotted the rank/location of our gene of interest against a line of common essential genes (red line in our plot) for that lineage. The red line indicates pan-essentiality^86^.

### Single-cell data analysis

Three public scRNA-seq datasets from primary prostate cancer were downloaded from GEO or a website provided by the author (GSE193337, GSE185344, www.prostatecellatlas.org)^86^. Using cell annotation from the Tuong et al. dataset as reference, luminal cells were annotated for the other two datasets with the label transfer method of Seurat^29^. Pseudo-bulk expression profiles^30^ were generated by summing counts from all cells annotated as luminal cells for each patient (tumor and normal samples separately). Normalization was achieved by computing normalization factors with the trimmed mean of M-values method^31^ and applying the cpm function from edgeR^32^. Boxplots of NSD2 and PCA3 expression were generated with ggpubr^33^ and paired Wilcoxon test was used to test the significance of the difference between benign and tumor (only patients with paired benign and tumor samples were included).

### Immunohistochemistry (IHC) and immunofluorescence (IF)

Immunohistochemistry (IHC) was performed on 4-μm-thick formalin-fixed, paraffin-embedded (FFPE) tissue sections using anti-NSD2 mouse monoclonal primary antibody (catalog no. ab75359, Abcam), anti-AR rabbit monoclonal primary antibody (catalog no. 760-4605, Roche-Ventana), and anti-CK8 rabbit monoclonal primary antibody (catalog no. ab53280, Abcam). Singleplex IHC was carried out on the Ventana ULTRA automated slide staining system (Roche-Ventana Medical Systems) using the OmniView Universal diaminobenzidine (DAB) detection kit (catalog no. 760-500, Roche-Ventana) and Hematoxylin II (catalog no. 790-2208, Roche-Ventana) for counterstain. Staining was evaluated under 100× and 200× magnification using a bright-field microscope.

Multiplexing immunofluorescence (IF) was performed consecutively as NSD2, CK8, and AR monoclonal antibodies described above with each antibody application and signal development kit followed by a denaturation cycle using high temperature and stringency wash to strip off the previous primary and secondary antibody detection system. The sequence of antibody application and the sequence of fluorophores were based on multiple denaturation and stripping optimization steps on contextual positive and negative control tissues. The consecutive IF detection system for the above-mentioned sequence of antibodies was developed by using the Discovery FITC kit (catalog no. 760-232, Roche-Ventana), Discovery Red 610 (catalog no. 760-245), and the Discovery Cy5 kit (catalog no. 760-238, Roche-Ventana) with DAPI (Prolong gold anti-fade, catalog no. P36931, Invitrogen/Fischer-Scientific). The multiplex staining was evaluated under confocal LMS900 system by Carl Zeiss.

#### Quantification of IF and IHC signal

**IF:** ImageJ was used to quantify the IF signal. Briefly, the color adjustment was done uniformly on all images being analyzed. Colors were converted to an RGB format, and a uniform color threshold was applied to all images. Using the particle measurement options, the integrated density values were obtained. A normalized fluorescence value was calculated and plotted in GraphPad Prism to normalize the signal from different sized nuclei. The normalized fluorescence value or corrected total cell fluorescence (CTCF) was calculated as^34^:

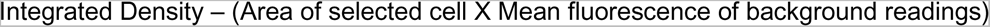

**IHC:** For NSD2 IHC quantification, the total number of immune-positive cells out of 100 (n/100) in four representative tumor cells and matched normal prostate epithelial cell regions was recorded and expressed as a percentage of positive cells for quantification purposes.

### Assessment of drug synergism

To determine the synergy between two drug treatments, cells were treated with increasing concentrations of either drug for 120 h, followed by the determination of viable cells using the CellTiter-Glo Luminescent Cell Viability Assay (Promega). The experiment was carried out in four biological replicates. The data were expressed as percentage inhibition relative to baseline, and the presence of synergy was determined by the Bliss method using the synergy finder R package.

### Animal procurement

Animal studies were approved by the Institutional Animal Care and Use Committee (IACUC) at the University of Pennsylvania and/or the University of Michigan. Animal use and care were in strict compliance with institutional guidelines, and all experiments conformed to the relevant regulatory standards by the universities. NOD SCID or NCI SCID/NCr athymic nude mice were obtained from the Jackson Laboratory (strain code: 005557) and Charles River (strain code: 561). All mice were housed in a pathogen-free animal barrier facility, and all *in vivo* experiments were initiated with male mice aged 5-8 weeks.

### Murine prostate tumor xenografts

Two million 22RV1 expressing Cas9, NSD2 KO, NSD2 rescue, and NSD2-dTAG cells were subcutaneously injected into the right flank or both dorsal flanks of 5–8-week-old male non-castrated and castrated NCI/NOD SCID/NCr mice (Charles River, strain code: 561, Jackson Laboratory, strain code: 005557) in 50% Matrigel (Corning, 354234). Castration surgery was performed by Charles River before receipt of the mice. The tumor measurement study ended at 60 days, and mice with tumors were sacrificed due to body condition or used for subsequent analysis. Tumor tissue was harvested and snap-frozen in liquid nitrogen for immunoblotting analysis. Mice that did not form tumors at this time continued to be monitored for tumor growth and body weight for first detection of tumor analysis up to 80 days. The chi-square- or t-test was used to evaluate the association of individual tumor characteristics with engraftment.

dTAGV-1 was formulated by dissolving into DMSO and further diluting with 10% solutol (Sigma): 0.9% sterile saline (Moltox) (w:v) with the final formulation containing 5% DMSO. IP injections were given at 25 mg/kg of body weight every 24 hours.

### Drug preparations: *in vitro* and *in vivo*

Enzalutamide (Selleckchem, S1250) and dTAG-13 (Sigma, SML2601) were dissolved and aliquoted in DMSO (Sigma, D2650). ARD61 was a kind gift from Dr. Shaomeng Wang. R1881 and UNC6934 were purchased from Sigma Aldrich, and JQ1 and EPZ-6438 were obtained from Selleck Chemicals. LLC0150 (NSD1/2) PROTAC was obtained in collaboration with Dr. Ke Ding at Shanghai Institute of Organic Chemistry, Chinese Academy of Sciences.

### Synthesis of LLC0150

General information: All chemicals and solvents were obtained from commercial suppliers and used without further purification. Purification was performed using combi-flash Nextgen300. All reactions were monitored by TLC, using silica gel plates with fluorescence F254 and UV light visualization. 1 H NMR spectra was recorded on a Varian Mercury Plus at 400 MHz, and 13C NMR spectra was recorded on a JEOL-ECZ-400S spectrometer at 100 MHz. Coupling constants (J) are expressed in hertz (Hz). Chemical shifts (δ) of NMR are reported in parts per million (ppm) units relative to internal control (TMS). Signal splitting patterns are described as singlet (s), doublet (d), triplet (t), quartet (q), multiplet (m), broad (br), or a combination thereof. The low resolution of ESI-MS was recorded on an Agilent-6120 and the high-resolution mass (resolution-70000) for compound was generated using Q-Exactive Plus orbitrap system, (Thermo Scientific) using electrospray ionization (ESI). Melting point was recorded in Stuart instrument, model smp30. Specific optical rotation was recorded in an Anton Paar MCP 5100 instrument. HPLC was recorded with a Waters 2696, and the column used was a YMC Triart C-18 EXRS (150*4.6) mm 5µm using 0.01M ammonium acetate in (Aq); Mobile phase-B: ACN 100%; Method -T/%B: 0/10, 2/10, 5/85, 13/85, 14/10, 15/10 method and flow rate: 1.0 ml/min. FT-IR was recorded using PerkinElmer Spectrum.

Abbreviations used: DMSO for dimethylsulfoxide, DIPEA for N, N-diisopropylethylamine, MeOH for methanol, DMF for N, N-dimethylformamide, HATU for 1-[bis(dimethylamino) methylene]-1H-1,2,3-triazolo[4,5-b]pyridinium 3-oxid hexafluorophosphate, DCM for dichloromethane, Pd(dppf)Cl2 for [1,1′-bis(diphenylphosphino)ferrocene]dichloropalladium(II)

Synthesis of *N*-cyclopropyl-*N*-(4-((4-((4-((2-(2,6-dioxopiperidin-3-yl)-1,3-dioxoisoindolin-5-yl)amino)butyl)carbamoyl)phenyl)carbamoyl)benzyl)-3-oxo-3,4-dihydro-2H-benzo[*b*][1,4]oxazine-7-carboxamide (LLC0150).

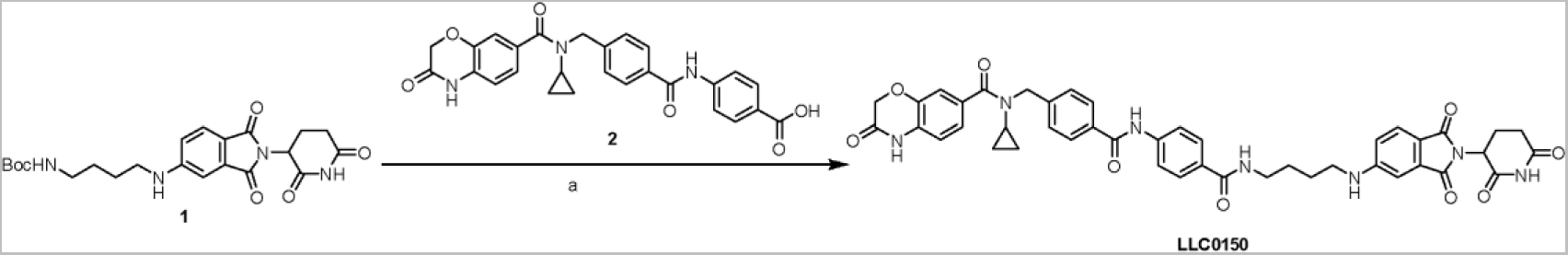

Reagents and conditions: (a) 1) trifluoroacetic acid (TFA), CH_2_Cl_2_, room temperature (rt), 2 h; 2) **2**, HATU, Et_3_N, DMF, rt, 3 h, 83% (two steps).

*tert*-butyl(4-((2-(2,6-dioxopiperidin-3-yl)-1,3-dioxoisoindolin-5-yl)amino)butyl)carbamate **1** (40.0 mg, 0.0900 mmol) was dissolved in 2.0 mL of CH_2_Cl_2_. To above solution was added TFA (0.5 mL) by syringe. The mixture was stirred at room temperature for two hours. After the reaction was complete, the solvent was removed by vacuum. The crude product was dissolved in a small amount of DMF (2.0 mL) and were added 4-(4-((*N*-cyclopropyl-3-oxo-3,4-dihydro-2H-benzo[*b*][1,4]oxazine-7-carboxamido)methyl)benzamido)benzoic acid **2** (48.1 mg, 0.0990 mmol), triethylamine (27.3 mg, 38 μL, 0.270 mmol), and HATU (51.3 mg, 0.135 mmol). The mixture was stirred at room temperature for three hours. The resulting mixture was purified by column chromatography to afford the title compound as a yellow solid (60.6 mg, 83% yield). ^1^H NMR (600 MHz, DMSO-*d*_6_) δ 11.05 (s, 1H), 10.88 (s, 1H), 10.43 (s, 1H), 8.40 (t, *J* = 5.6 Hz, 1H), 7.96 (d, *J* = 8.2 Hz, 2H), 7.88 – 7.82 (m, 4H), 7.55 (d, *J* = 8.4 Hz, 1H), 7.46 (d, *J* = 7.7 Hz, 2H), 7.22 – 7.13 (m, 3H), 6.98 – 6.95 (m, 1H), 6.93 (d, *J* = 8.1 Hz, 1H), 6.86 (dd, *J* = 8.4, 1.9 Hz, 1H), 5.02 (dd, *J* = 12.8, 5.5 Hz, 1H), 4.72 (s, 2H), 4.62 (s, 2H), 3.30 (d, *J* = 6.2 Hz, 2H), 3.23 – 3.19 (m, 2H), 2.90 – 2.84 (m, 1H), 2.83 – 2.77 (m, 1H), 2.62 – 2.51 (m, 2H), 2.02 – 1.96 (m, 1H), 1.67 – 1.59 (m, 4H), 0.58 – 0.52 (m, 2H), 0.50 – 0.45 (m, 2H); ^13^C NMR (151 MHz, DMSO-*d*_6_) δ 172.82, 170.19, 167.71, 167.15, 165.64, 165.57, 164.85, 154.46, 142.49, 142.35, 141.70, 134.22, 133.42, 131.79, 129.49, 128.39, 128.06, 127.83, 127.21, 125.10, 121.87, 119.34, 115.81, 115.36, 115.16, 66.74, 48.61, 42.23, 40.06, 38.77, 30.98, 26.82, 25.75, 22.23, 9.55; HRMS (m/z): [M + Na]+ calcd for C_44_H_41_N_7_O_9_Na+ 834.2858, found 834.2861; HPLC purity: 97.92%.

### ^1^H NMR of (LLC0150)

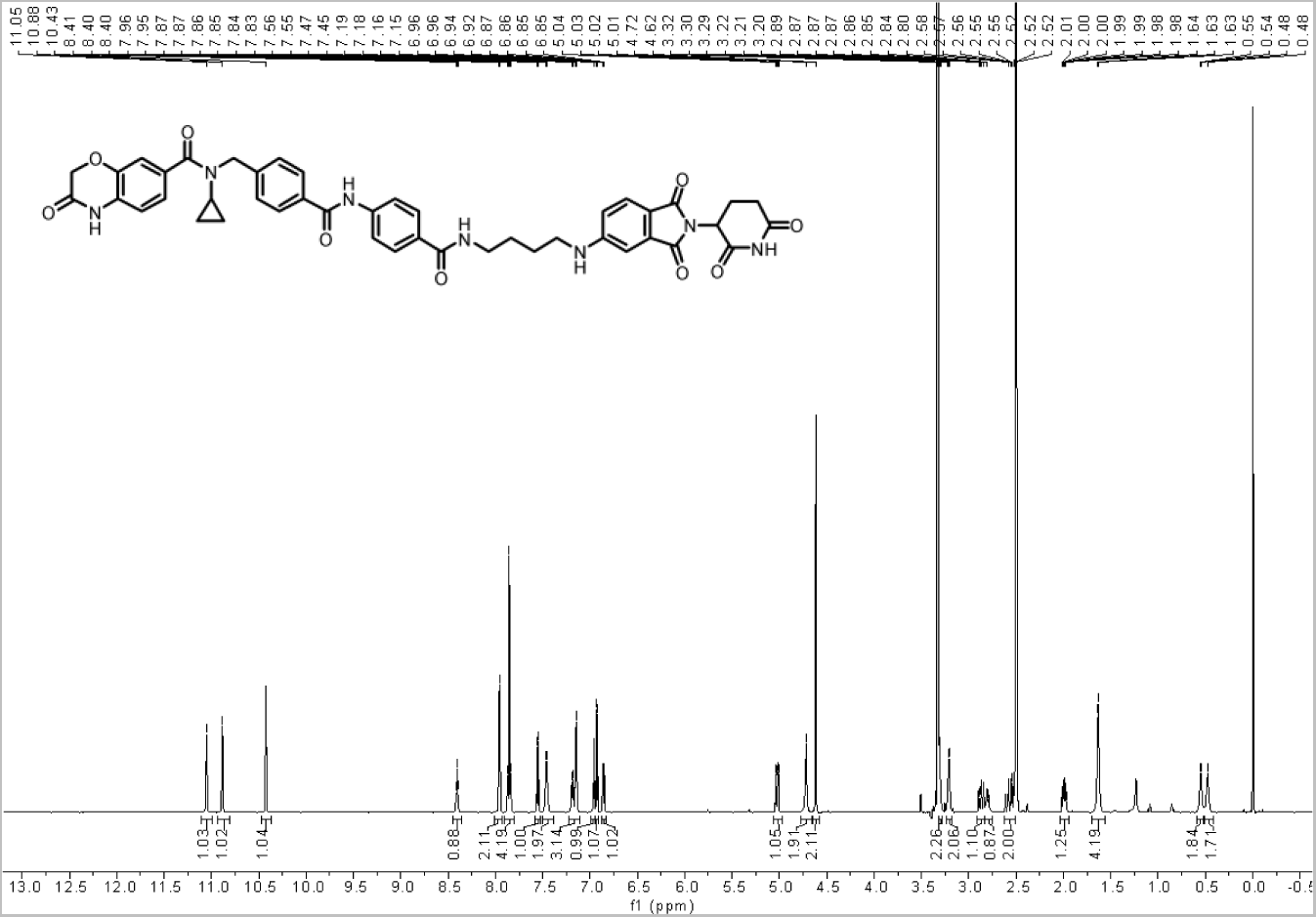

### ^13^C NMR of LLC0150

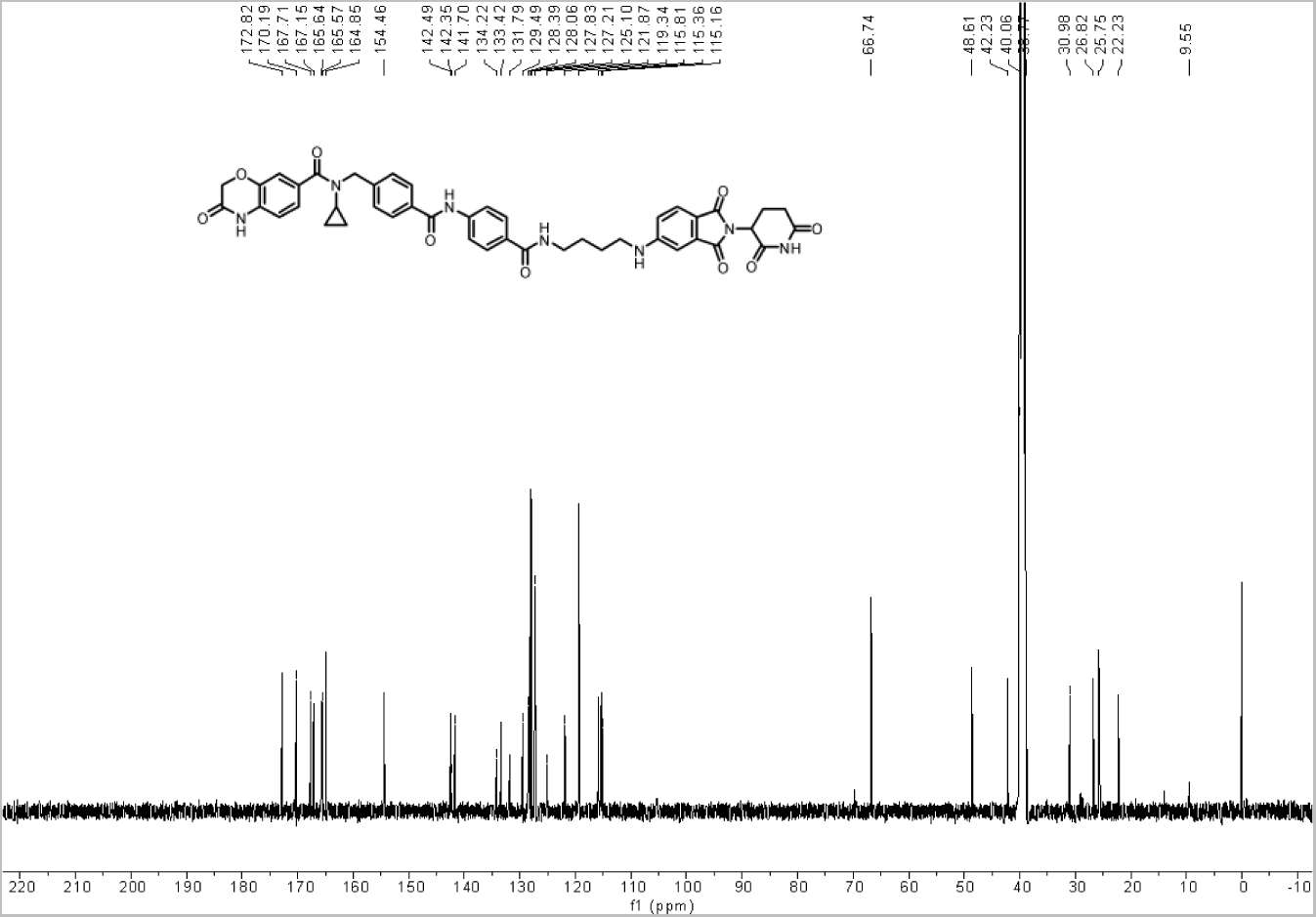

### Plasmids and lentivirus

The pMSCV-HA-NSD2-PH WT and pMSCV-HA-NSD2-PH-W236A, pMSCV-HA-NSD2-PH-F266A, pMSCV-HA-NSD2-PH-W236A/F266A, pMSCV-HA-HMGa, pMSCV-HA-HMGa-F463A, pMSCV-HA-HMGa-W491A, and pMSCV-HA-HMGa-Y502A constructs were created through site-directed mutagenesis. Site-directed mutagenesis was performed using the QuikChange II XL site-directed Mutagenesis Kit (Agilent Technologies, 200521) as per the manufacturer’s protocol. All mutations were verified by Sanger sequencing. The following primers from Integrated DNA Technologies (IDT) were used for site-directed mutagenesis and cloning:

#### Site-directed mutagenesis

pMSCV-NSD2-PH-W236A: F-5’-AGA AAC CAT GCA AGG CGC CCA AGG GTA ACC CGA C-3’, R-5’-GTC GGG TTA CCC TTG GGC GCC TTG CAT GGT TTC T-3’; pMSCV-NSD2-PH-F266A: F-5’-CTG GGG CGT CAC CAA AGG CCT GTA CGT GAT ACT GGC-3’, R-5’-GCC AGT ATC ACG TAC AGG CCT TTG GTG ACG CCC CAG-3’; pMSCV-NSD2-HMGa-F463A: F-5’-CCAT CCC TGT GTT TTT GAC AGG CGA CCA AAA ACT GGG ATG CTG-3’, R-5’-CAG CAT CCC AGT TTT TGG TCG CCT GTC AAA AAC ACA GGG ATG-3’; pMSCV-NSD2-HMGa-W491A: F-5’-CTC ACT CAG CAG ACT CGC CTG TGA CCT GAG CAG C-3’, R-5’-GCT GCT CAG GTC ACA GGC GAG TCT GCT GAG TGA G-3’; pMSCV-NSD2-HMGa-Y502A:F-5-’GGG CAA ACT TGG TGT TGG CGC GTG CTC TCT GCT TC-3’, R-5’-AGA AGC AGA GAG CAC GCG CCA ACA CCA AGT TTG CCC-3’

#### Cloning

pMSCV-HA-NSD2-Fl: F-5’AAG GAT CCG AAT TTA GCA TCA AG-3’, R-5’-TTC TCG AGC TAT TTG CCC TCT GTG AC-3’; pMSCV-HA-NSD2-PH: F-5’-AAG GAT CCG AAT TTA GCA TCA AG-3’, R-5’-TTC TCG AGT TAT CGC TCA GAC TTT TTG GA-3’; pMSCV-HA-NSD2-PH: F-5’-CAA ATG TTG CTT GTC TGG TG-3’, R-5’-GTC AGT CGA GTG CAC AGT TT-3’; pMSCV-HA-NSD2-HMGa: F-5’-TTC TCG AGT TAA TGG AGG AGC GGA AAG CCA AGT T-3’. R-5’-TTC TCG AGT TAT CGC TCA GAC TTT TTG GA-3’; pMSCV-NSD2-HMGa-Y502A:F-5’GGG CAA ACT TGG TGT TGG CGC GTG CTC TCT GCT TC-3’, R-5’AGA AGC AGA GAG CAC GCG CCA ACA CCA AGT TTG CCC-3’;pMSCV-HA-NSD2-PWWP1a: F-5’-AAG GAT CCG AAT TTA GCA TCA AGC AG-3’, R-5’-TTC TCG AGT TAC CTT CTC TTC ATG GGA AT-3’;pJS60-FKBP5-HA-NSD2-Fl: F-5’-GGTGTCGTGACGCGGATGGAATTTAGCATCAAGCAGAGTCC-3’,R-5’-TTCCACCTGCACTCCTTTGCCCTCTGTGACTCTCCG-3’;pJS60:F-5’-CCGCGTCACGACACCT-3’,R-5’-GGAGTGCAGGTGGAAACCA-3’;pCDNA4c:F-5’-GCG GCC GCT CGA GTC-3’, R-5’-GGA TCC TGG TAC CTT ATC GTC AT-3’;NSD2-HMGa: F-5’-AAG GTA CCA GGA TCC ATG CGG AGG GCC AAA CT-3’, R-5’TCG AGC GGC CGC CTA TCG CTC AGA CTT TTT GGA TGA GAC AG-3’

Other plasmids used in this study were pMSCV-HA-NSD2-FL, pJS60-FKBP5-HA-NSD2-FL, pCDNA4c-HMGa, pFN21K-Halo-AR-FL, pFN21K-Halo-AR-NTD, pFN21K-Halo-AR-DBD, and pFN21K-Halo-AR-HBD. The plasmids were transfected into HEK293-T or LNCaP cells using Lipofectamine 2000 (Invitrogen) as per the manufacturer’s protocol. The cells were then harvested 48/72 h post-transfection and subjected to immunoprecipitation or immunoblotting or used for lentivirus production. Lentiviral particles were generated in the lab at the University of Pennsylvania or in collaboration with the Vector Core at the University of Michigan. Following lentiviral production, viral particles were incubated on cells in penicillin/streptomycin-free media for 24 h, followed by media replacement with normal media. Overexpression and knockdown of proteins was confirmed by immunoblotting.

## Data and materials availability

All data are available in the manuscript or the supplementary information. All materials are available from the authors upon request. All raw next-generation sequencing data, including ChIP-seq and RNA-seq, generated in this study are deposited in the NCBI Gene Expression Omnibus (GEO) repository (accession number: GSE242737).

