## Supplementary figures and images for "NSD2 is a requisite subunit of the AR/FOXA1 neo-enhanceosome in promoting prostate tumorigenesis"

### Supplemental Figures 1 - 7

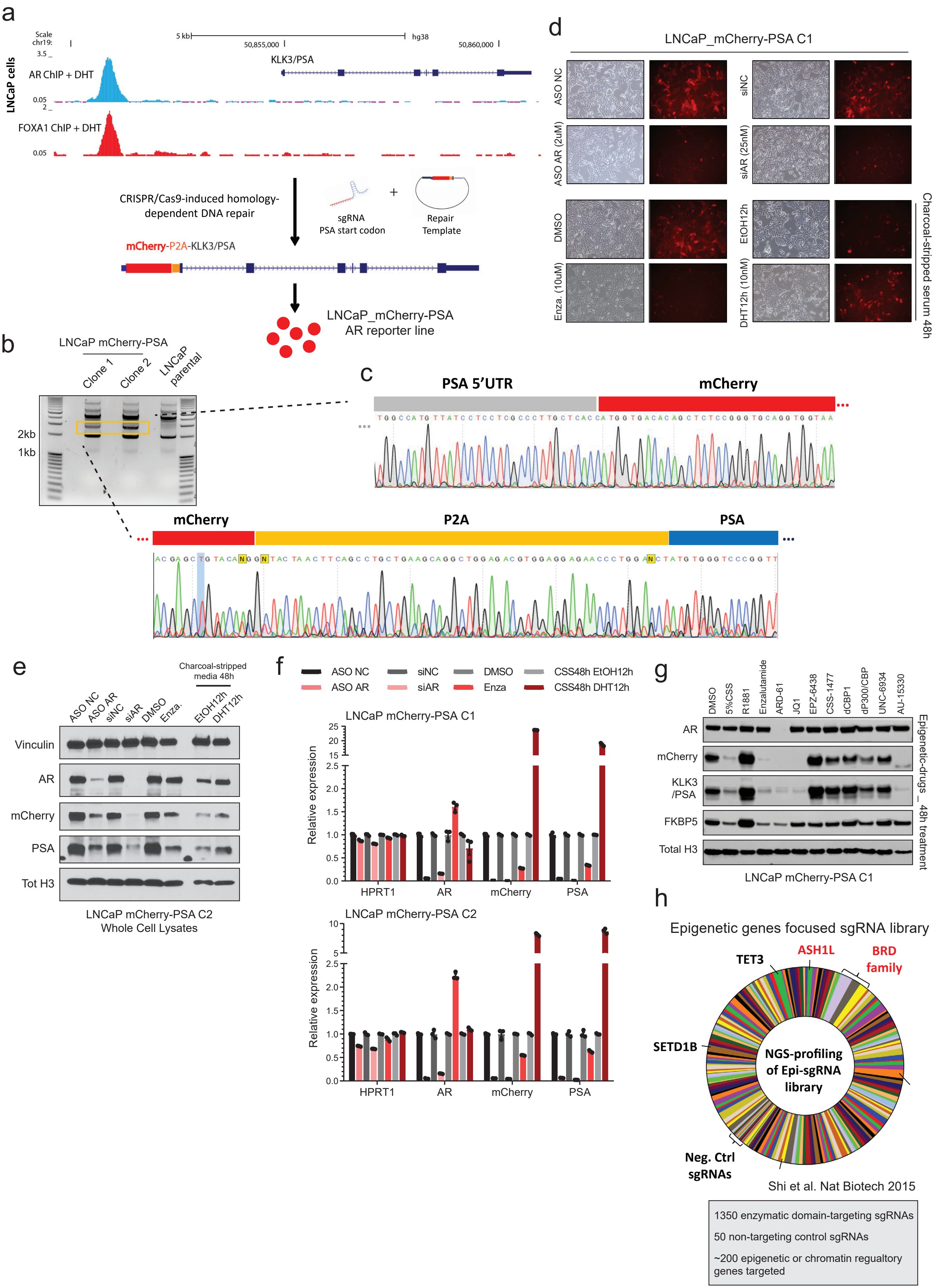

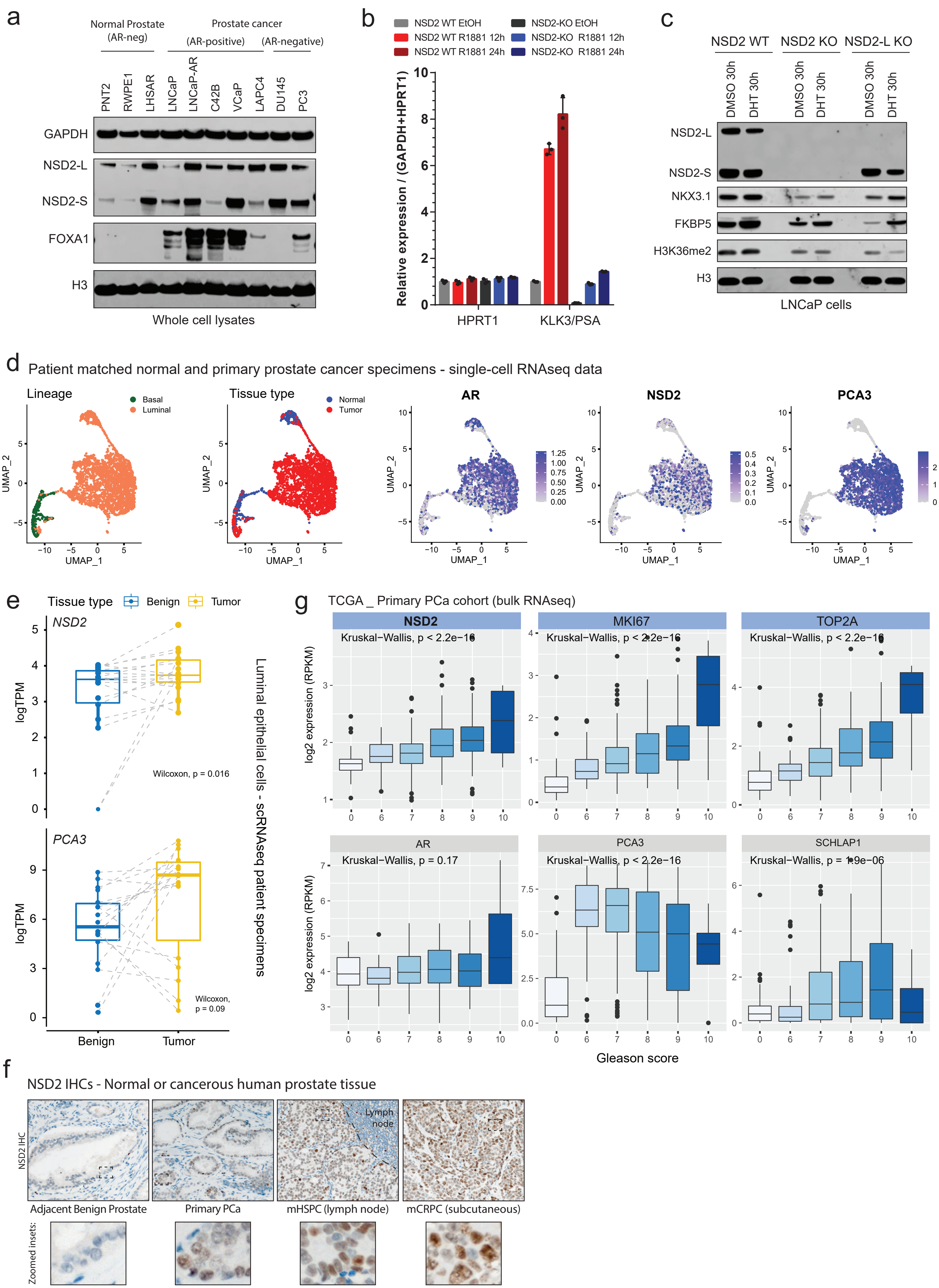

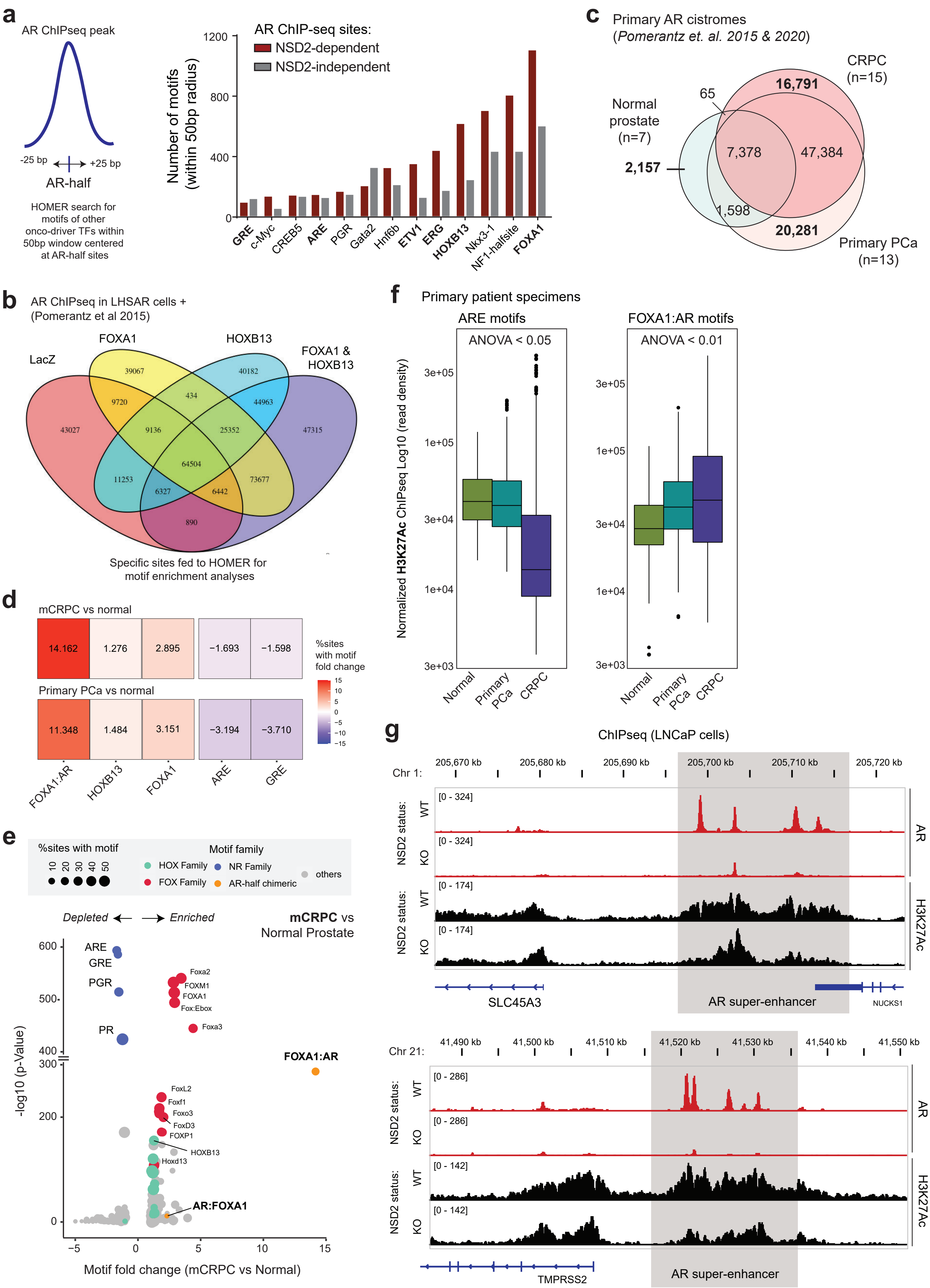

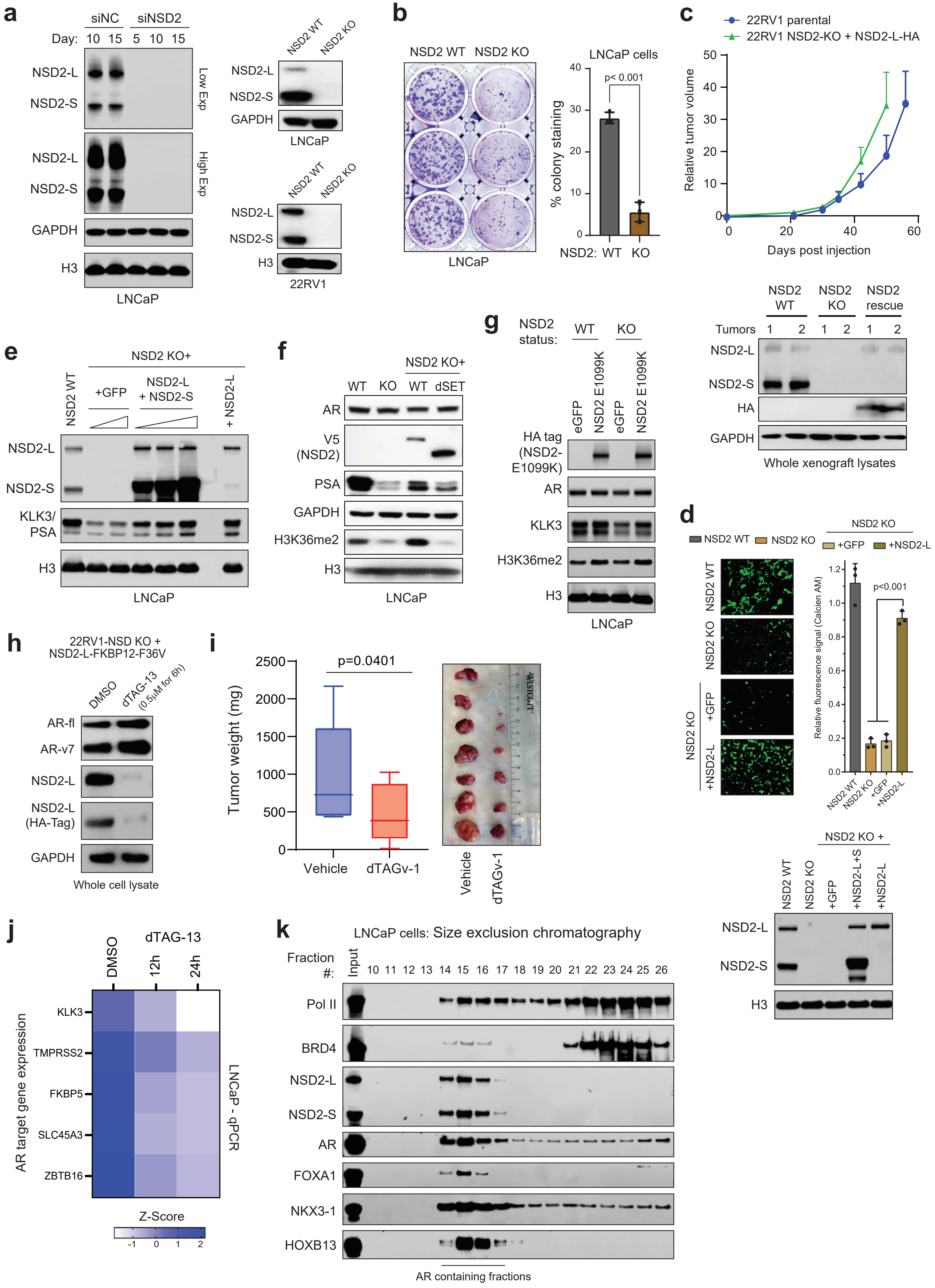

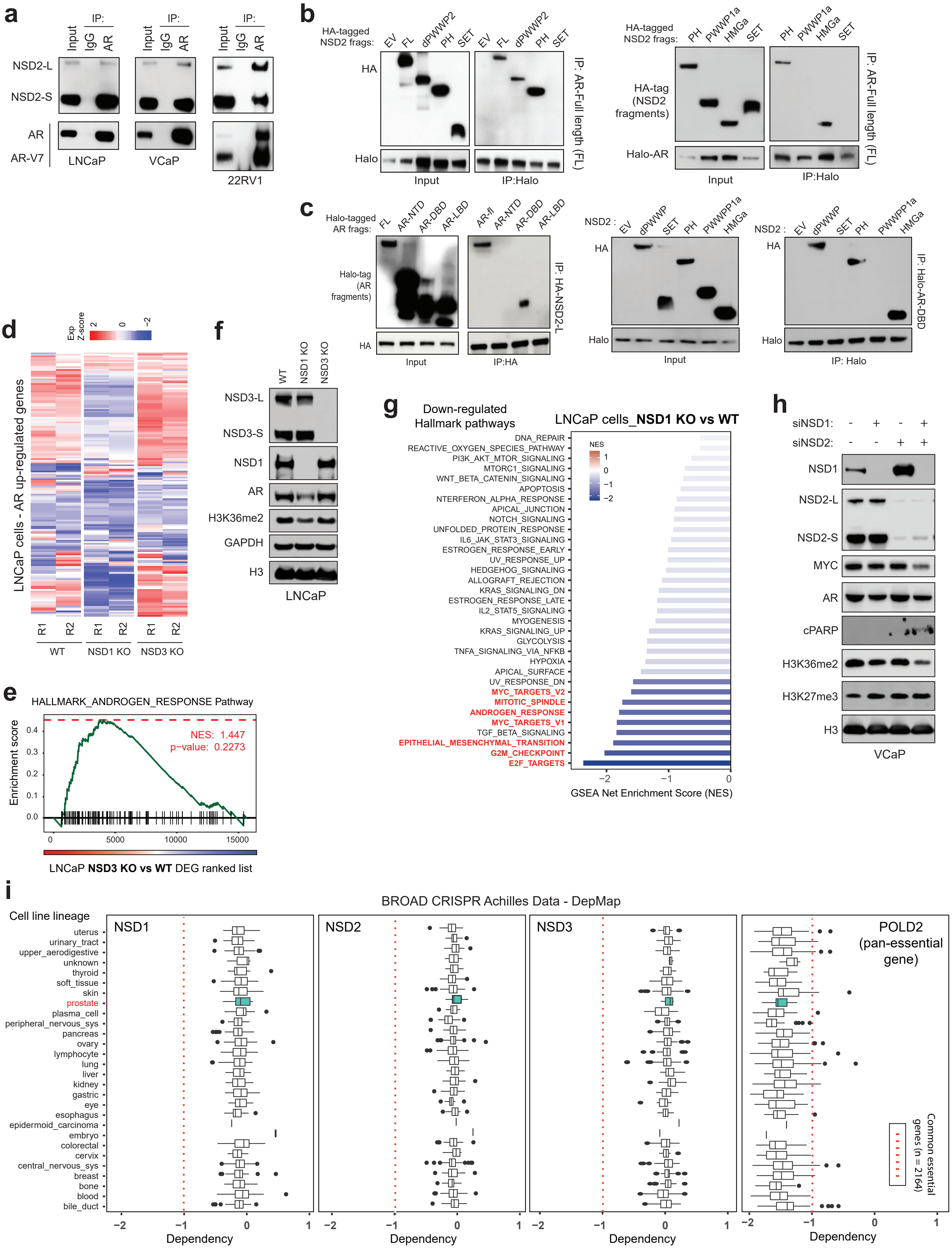

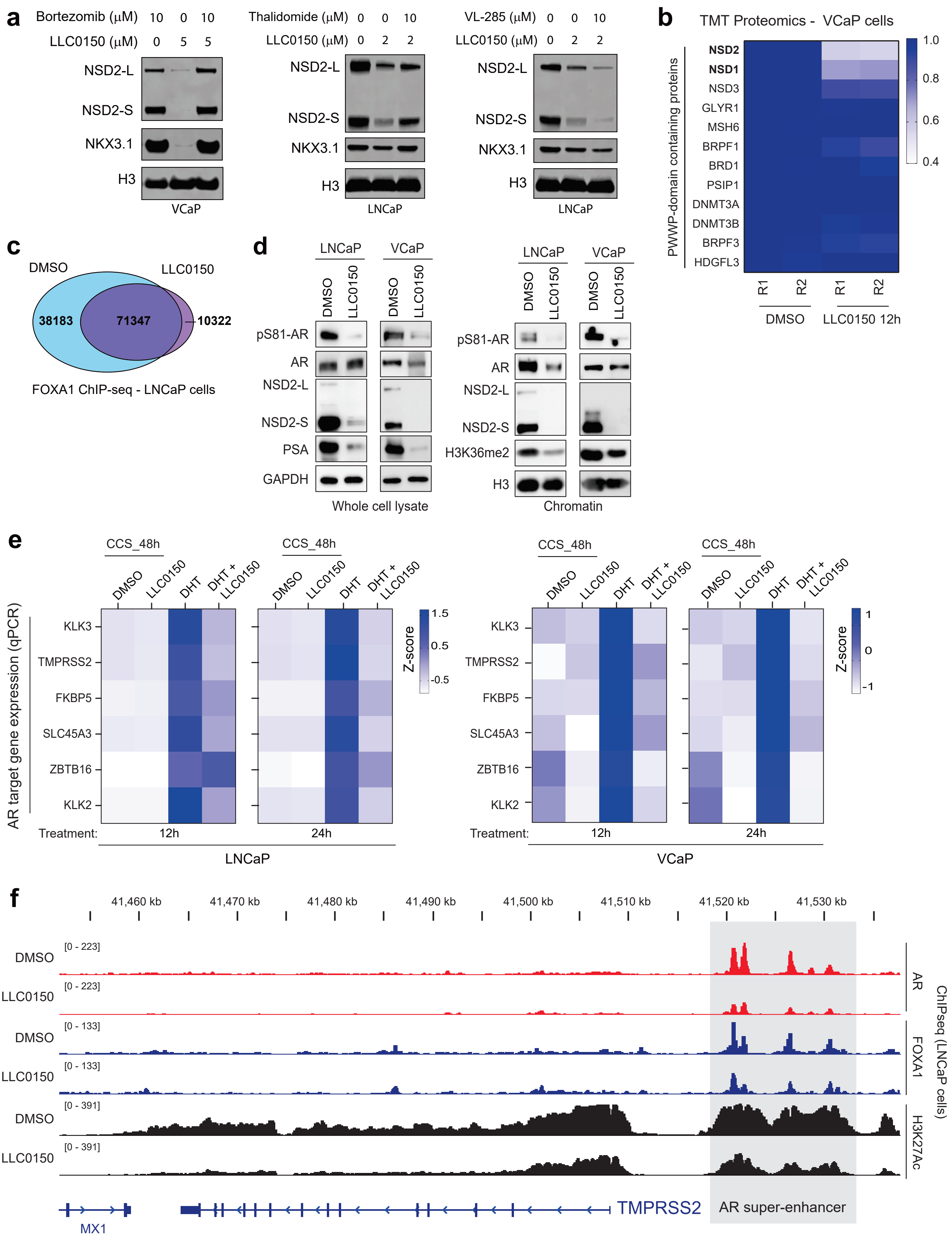

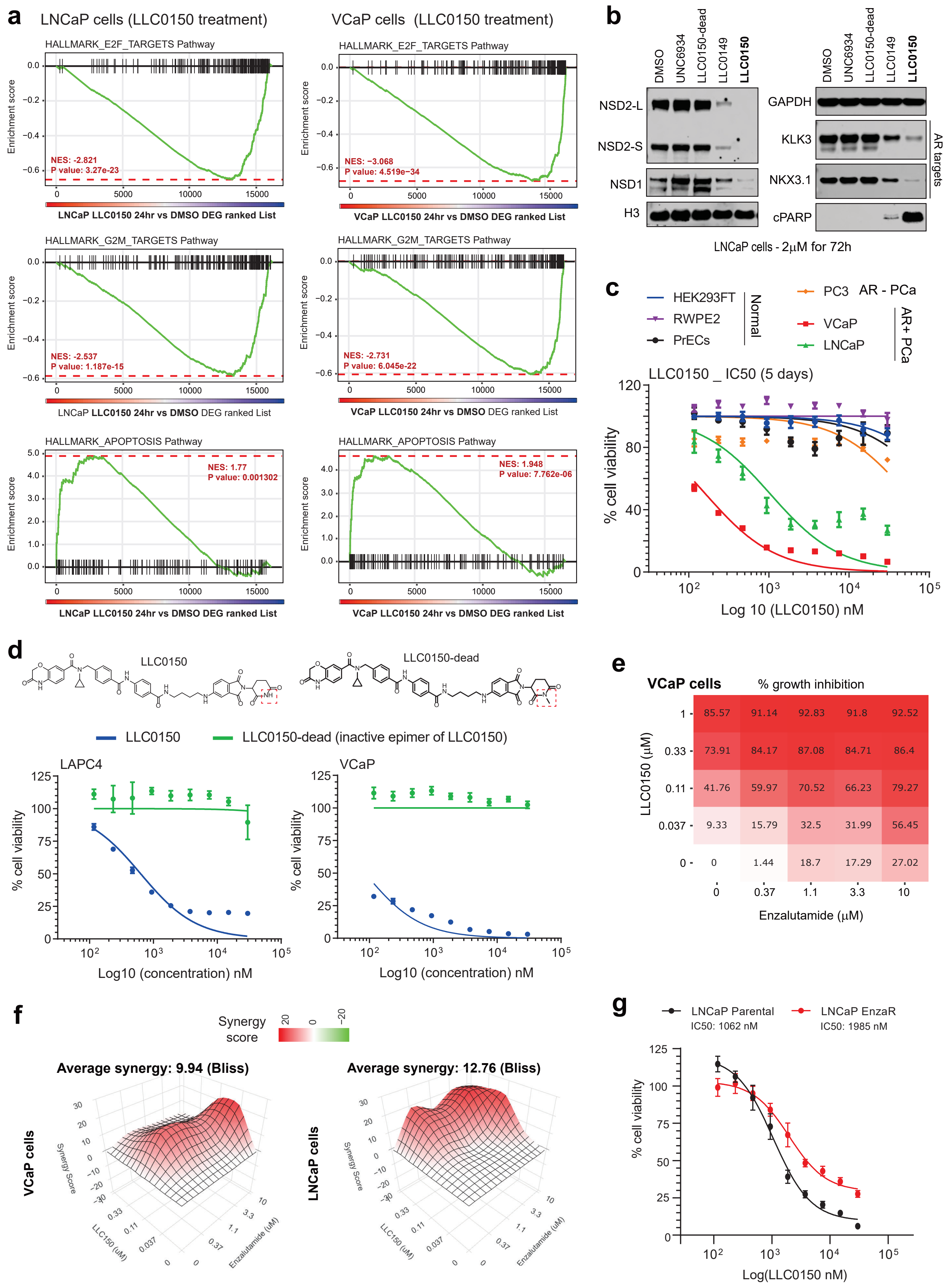
